# Cofilin loss in *Drosophila* contributes to myopathy through defective sarcomerogenesis and aggregate formation during muscle growth

**DOI:** 10.1101/825448

**Authors:** Mridula Balakrishnan, Shannon F. Yu, Samantha M. Chin, David B. Soffar, Stefanie E. Windner, Bruce L. Goode, Mary K. Baylies

## Abstract

Sarcomeres, the fundamental contractile units of muscles, are conserved structures composed of actin thin filaments and myosin thick filaments. How sarcomeres are formed and maintained is not well understood. Here, we show that knockdown of *Drosophila* Cofilin (*DmCFL)*, an actin depolymerizing factor, leads to the progressive disruption of sarcomere structure and muscle function *in vivo*. Loss of *DmCFL* also results in the formation of sarcomeric protein aggregates and impairs sarcomere addition during growth. Strikingly, activation of the proteasome delayed muscle deterioration in our model. Further, we investigate how a point mutation in CFL2 that causes nemaline myopathy (NM) in humans, affects CFL function and leads to the muscle phenotypes observed *in vivo*. Our data provide significant insights to the role of CFLs during sarcomere formation as well as mechanistic implications for disease progression in NM patients.

## INTRODUCTION

Skeletal muscle cells, or myofibers, are a highly organized cells that are necessary for locomotion. Each myofiber contains many linear myofibrils, which span the entire length of the cell. Each myofibril, in turn, is composed of a repeated array of sarcomeres, the fundamental contractile units of the muscle cell. The coordinated contraction of all sarcomeres along the myofibril shortens the entire myofiber and produces mechanical force. Sarcomeres are conserved in both structure and function amongst metazoans and have been studied for decades. However, much remains unknown as to how sarcomeres are assembled and incorporated into myofibrils during muscle development and growth.

Sarcomeres consist of three structural elements: Z-discs, thin filaments, and thick filaments (Figure 1A). The Z-discs define the lateral sarcomere boundaries and are composed of electron dense proteins, including *α*-actinin and ZASP/Cypher. The thin filaments are anchored at the Z-discs and extend towards the middle of the sarcomere (M-line). Thin filaments are comprised primarily of actin and actin binding proteins, such as nebulin and the troponin-tropomyosin complex. The actin filaments are capped at their pointed (minus) ends by tropomodulin (Tmod) and at their barbed (plus) ends by CapZ, which also anchors the thin filaments at the Z-disc. The thick filaments are at the center of the sarcomere and are composed primarily of myosin and myosin associated proteins. Thick filaments interact with thin filaments, and are crosslinked at the M-line, which is composed of proteins like obscurin (Henderson et al., 2017). While many conserved sarcomeric proteins have been identified, their incorporation into and function within the sarcomere are not completely understood. Moreover, it is still unclear as to how sarcomere size is regulated and maintained during muscle homeostasis. Understanding these fundamental areas of muscle biology is crucial, as mutations in sarcomeric proteins have been implicated in several muscle diseases.

**Figure 1:**
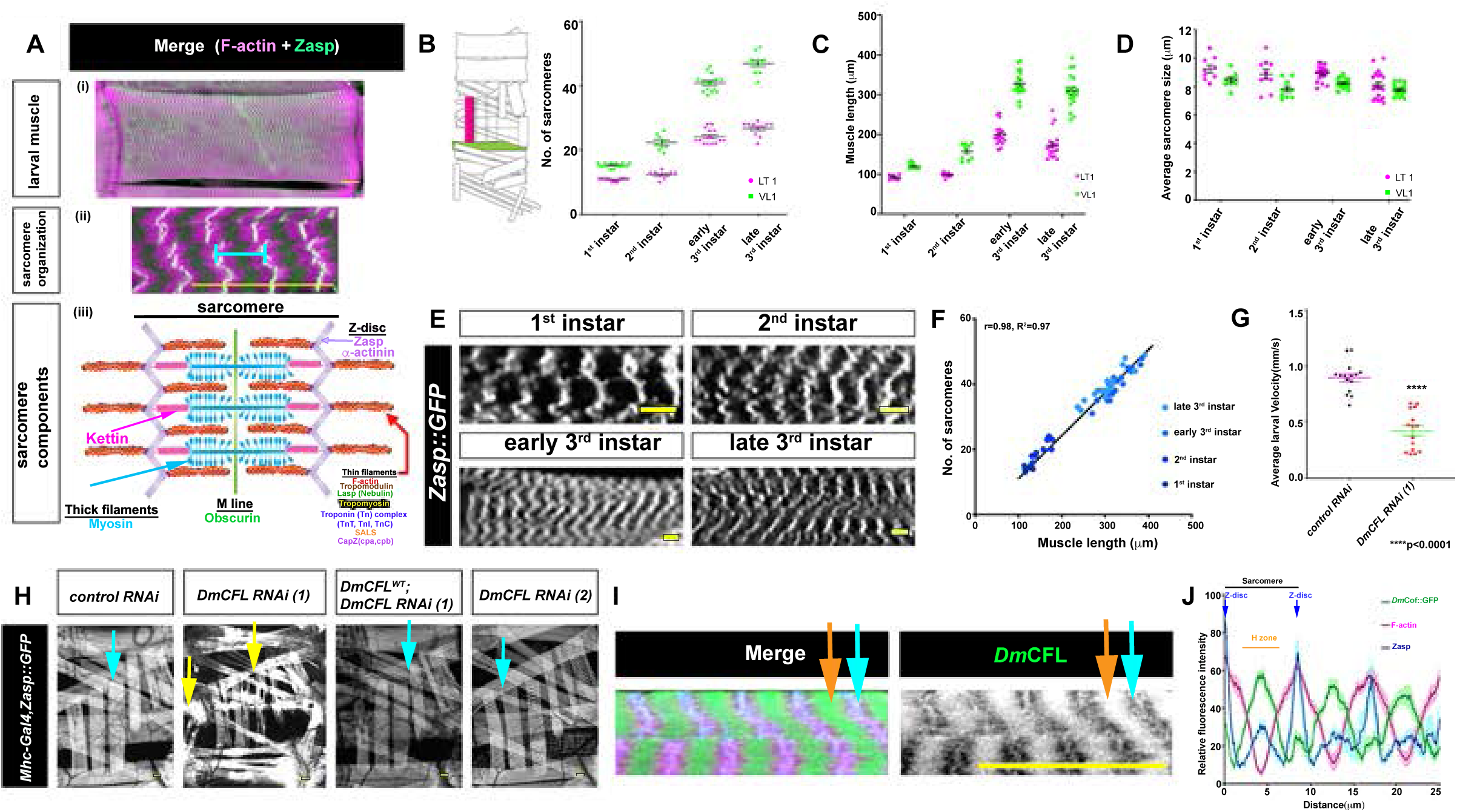
Muscle-specific knockdown of *Dm*CFL affects muscle structure and function. **(A)** (i,ii) *Drosophila* larval muscle at the 3^rd^ instar stage showing sarcomere organization [Z-disc (*α*-Zasp, green) and F-actin (phalloidin, magenta)]. Blue line indicates one sarcomere in (ii). (iii) Schematic depicting a single sarcomere with its structural components: Z-discs, thin filaments, and thick filaments. The *Drosophila* proteins analyzed in this study and their location within the sarcomere are marked in corresponding color. **(B)** (Left) Diagram of one hemisegment of the 3^rd^ instar larval musculature. Two different *Drosophila* larval muscles, LT1 (magenta) and VL1 (green) are highlighted. (Right) Graph depicting sarcomere number in LT1 (magenta) and VL1 (green) at each larval instar. N=15 muscles from at least 5 *control RNAi* larvae at each instar. **(C)** Graph depicting muscle length in LT1 (magenta) and VL1 (green) muscles at each larval instar. N=15 muscles from at least 5 *control RNAi* larvae at each instar. **(D)** Quantification of sarcomere length in LT1 (magenta) and VL1 muscles (green) at each larval instar stage (mean±SEM). N=15 muscles from at least 5 control larvae at each instar. **(E)** Sarcomeres at the different larval instar stages, as revealed by Zasp-GFP expression that labels Z-lines (greyscale). Scale Bars 8 µm. **(F)** Relationship between VL muscle cell length and number of sarcomeres at each larval stage. Line indicates linear scaling relationship (Pearson’s correlation coefficient (r) and R^2^ are indicated). **(G)** Average velocities of early 3^rd^ instar larvae expressing *Mhc-Gal4* with either *control RNAi* or *DmCFL RNAi (1)* (mean±SEM). Data are representative of one experiment of each genotype. Experiment was repeated 2 times with a N=15 larvae of each genotype with similar results. P values calculated by Unpaired *t*-test. **(H)** Larval muscles at the 3^rd^ instar stage expressing Zasp-GFP (greyscale), and *Dmef2-Gal4* to drive different RNAi’s as indicated. Several individual muscles of one hemisegment are shown. Muscles expressing *DmCFL RNAi (1)* show significant defects (Zasp aggregates, yellow arrows). Expression of *DmCFL^WT^* in this RNAi background rescues this phenotype. Dm*CFL RNAi (1)* had no effect on muscle structure using this driver. Blue arrows denote examples of wildtype sarcomere pattern; yellow arrows denote examples of disorganized sarcomeres and Zasp aggregates. **(I)** Localization of *Dm*CFL-GFP (α-GFP, green (left panel); greyscale (right panel)), F-actin (phalloidin, magenta), and Zasp proteins (α-Zasp, blue) in larval sarcomeres. Orange arrow: H zone; blue arrow: Z-disc. **(J)** Relative average fluorescence intensities of these proteins across the length of 4 sarcomeres (average ± SEM). *Dm*CFL-GFP localizes to the Z-discs of the sarcomere as well as the F-actin free region, i.e. the H-zone. N = 10 measurements, from at least 5 different muscles from 5 larvae. For endogenous *Dm*CFL-GFP pattern see Supplementary Figure 1J. Scale bars: 25 µm (A,H, I), 8 µm (E).

A skeletal muscle disease with links to sarcomere structure and sarcomeric proteins is nemaline myopathy (NM). NM has an incidence of 1 in 50,000 births (Maggi et al., 2013) and involves a range of clinical symptoms, including difficulties in breathing and swallowing, hypotonia, delayed motor development, and atrophied muscle fibers (Malfatti and Romero, 2016; Sewry et al., 2019; Wallgren-Pettersson et al., 1999; 2004). Interestingly, not all skeletal muscles are affected equally in NM patients (Malfatti and Romero, 2016). A characteristic feature of affected muscles is the presence of electron dense protein aggregates within the muscle fibers, called nemaline (rod) bodies. These aggregates are composed of sarcomeric proteins, such as *α*-actinin(Cassandrini et al., 2017; Jockusch et al., 1980; Sewry et al., 2019; Wallgren-Pettersson et al., 1995) and *α*-actin (Cassandrini et al., 2017; Yamaguchi et al., 1982), and are thought to be extensions of the Z-disc (Romero et al., 2013). While nemaline bodies are the key diagnostic feature of NM, there is no correlation between the number of nemaline bodies and the severity of the disease (Malfatti and Romero, 2016). Mutations in at least 12 genes, many that encode sarcomeric proteins, have been implicated in NM (Gupta and Beggs, 2014; Kondo et al., 2012; Malfatti and Romero, 2016; Miyatake et al., 2017; Nilipour et al., 2018; Sandaradura et al., 2018; Sewry et al., 2019). Due to a scarcity of patient samples, it remains unclear how mutations in individual genes contribute to the development and progression of NM, particularly the mechanisms that result in the formation of nemaline bodies and muscle weakness.

One of the genes mutated in NM patients is *CFL2*, which encodes a member of the actin depolymerizing factor (ADF)/Cofilin family of proteins (Agrawal et al., 2007; Ockeloen et al., 2012; Ong et al., 2014). Humans possess 3 ADF/Cofilin family members: ADF/*DSTN* (*Destrin*), *CFL1* (*Cofilin-1*), and *CFL2* (*Cofilin-2*). *CFL2* is the primary member of this family that is expressed in mature skeletal muscle, and its loss results in aberrant skeletal muscle structure and early lethality (Agrawal et al., 2012; Ockeloen et al., 2012). Like other members of this family, Cofilin-2 binds to globular actin monomers (G-actin) to inhibit nucleotide exchange (ADP to ATP), and to filamentous actin (F-actin) to promote their fragmentation and disassembly into monomers (Vartiainen et al., 2002). *CFL2* directly severs actin filaments (Chin et al., 2016) and is thereby thought to have evolved to regulate thin filament length in the sarcomere (Kremneva et al., 2014a). While CFL2’s structure and function have been identified, how mutations in CFL2 contribute to the pathogenesis of NM is not well understood.

An ongoing *Drosophila* screen in our lab identified *twinstar* (*tsr*) as a putative regulator of muscle formation. Tsr is a *Drosophila* homolog of the Actin Depolymerizing Factor (ADF)/Cofilin family of proteins and has the highest sequence homology to human Cofilin-2 (*CFL2*; (Edwards et al., 1994; Gunsalus et al., 1995). Tsr (*Dm*CFL) has been shown to be involved in cytokinesis, axonal growth, and in planar cell polarity pathway (Blair et al., 2006; Gunsalus et al., 1995; Ng and Luo, 2004). Here we report a new role of *Dm*CFL in regulating both sarcomere and myofiber structure in the *Drosophila* larval musculature. Our data suggest that muscle-specific knockdown of *Dm*CFL impairs the addition of new sarcomeres during muscle growth, which results in a progressive loss of sarcomere and muscle fiber integrity. We observe the formation of protein aggregates within affected muscle fibers and directly link progressive muscle deterioration to deficits in locomotion behavior. Further, we demonstrate that increasing the activity of the proteasome improves muscle structure and function in *Dm*CFL-knockdown muscles. Finally, we demonstrate functional similarities between *Drosophila* and human Cofilin proteins through *in vitro* and *in vivo* analyses and model a specific mutation found in a human NM patient. Together, our data indicate that the first overt morphological change in the *Dm*CFL-knockdown muscle cells is the accumulation of F-actin at sites of active sarcomerogenesis, which leads to the recruitment of actin capping proteins from existing sarcomeres, myofibril degeneration, and muscle weakness. Our data provide significant insights to the role of CFL2 during muscle growth as well as mechanistic implications for disease progression in NM patients.

## RESULTS

### Muscle specific knockdown of *Dm*CFL disrupts muscle structure and function

The *Drosophila* larval musculature, which is established in the embryo, consists of a repeated pattern of hemisegments along the body wall, each of which consists of 30 unique muscles of different shapes, sizes, and orientations. Each larval muscle is a single cell or fiber, enabling easy visualization of the muscle fiber structure (Figure 1A; Bate, 1990). Individual myofibers as well as their sarcomeric organization can be easily visualized using an array of protein traps and fluorescent constructs which enable imaging of both live and fixed muscles. The larva, upon hatching, progresses through 3 developmental stages, known as instars (5 days at 25°C), during which the muscles grow 25-40-fold (Demontis and Perrimon, 2009). A study from the 1950’s showed that *Drosophila* larval muscles growth is fueled by increasing sarcomere number (HAAS, 1950). To confirm this using modern techniques, we quantified the number of sarcomeres in two different muscles, using a Zasp-GFP protein trap (Zasp::GFP) which marks the Z-disc of sarcomeres. We observed that both sarcomere number and muscle length increased in both muscles through the instars, while sarcomere size remained constant (Figures 1B-1E). In accordance with a consistent sarcomere size, quantification of both cell length and sarcomere number throughout larval development revealed a linear scaling relationship of those two parameters (r=0.98; Figure 1F). These data suggested the *Drosophila* larval musculature is an excellent system to study sarcomere assembly and addition, as well as maintenance.

To investigate the role of *Dm*CFL (Tsr) in the larval musculature, we used the *Gal4/UAS* system (Brand and Perrimon, 1993) to manipulate *Dm*CFL specifically in the muscle cells. Using an early embryonic muscle driver, *Dmef2-Gal4,* we expressed two different *UAS tsr RNAi* constructs, referred to as *UAS DmCFL RNAi (1)* and *UAS DmCFL RNAi (2)* (S1A). Expressing either RNAi resulted in a severe disruption of muscle fiber structure and sarcomere organization, and early death at the 2^nd^ instar stage. Expression of *DmCFL^WT^,* together with either of the *DmCFL RNAi* constructs, rescued both muscle and sarcomere structure as well as viability (S1B-S1D). Together, these experiments showed that targeted knockdown of *DmCFL* using a strong, early embryonic driver caused severe muscle phenotypes; however, the early larval death impeded detailed analyses of *Dm*CFL’s role in sarcomere organization and in muscle development and function.

To bypass this early larval lethality, we expressed both *DmCFL RNAi* constructs with the muscle Gal4 line, *Mhc-Gal4,* that drives expression later in embryogenesis and at lower levels relative to *Dmef2-Gal4* (Kaya-Çopur and Schnorrer, 2019; Viswanathan et al., 2015; Zhang and Bernstein, 2001). While the expression of *DmCFL RNAi (2*) with *Mhc-Gal4* did not cause any muscle phenotypes or viability issues as a result of insufficient *Dm*CFL knockdown, expression of *DmCFL RNAi (1)* resulted in organism death after the larval stages (Figures 1G and 1H, S1E-S1G). At the 3^rd^ instar stage, *DmCFL RNAi* (1) knockdown larvae showed slower locomotion and disrupted muscle structure with the formation of Zasp aggregates within the muscle fibers (Figures 1G and 1H). Expression of *DmCFL^WT^* together with *DmCFL RNAi (1)* rescued viability and muscle structure, while expression of *2x-GFP* together with *DmCFL RNAi (1)* failed to improve viability (Figure 1H, S1E and S1F). These data confirmed that the specificity of the knockdown and rescue phenotypes that we observed in the larval muscles. Consistent with an effect of *Dm*CFL’s loss on sarcomere structure, we found that *Dm*CFL’s localization is restricted to the Z-disc and H-zone of sarcomeres in the 3^rd^ instar muscle (Figures 1I and 1J, S1J). This expression was lost upon *Dm*CFL knockdown (S1G); immunoblotting of whole 3^rd^ instar larvae also revealed reduction of *Dm*CFL levels to 60% of control levels (S1K).

Together these data showed that using a specific combination of genetic tools (*Mhc-Gal4*, *UAS DmCFL RNAi (1))* we developed a *Drosophila* model for the detailed analysis of sarcomeric organization in *Dm*CFL-knockdown muscles during a period of muscle growth.

### *Dm*CFL knockdown results in three classes of muscle phenotypes with varying protein aggregate composition

Our initial analyses showed that muscle-specific knockdown of *Dm*CFL leads to disruption of muscle fiber structure and sarcomere organization, with a concomitant reduction in muscle function and viability. To further investigate the structural changes within the muscle fibers, we examined the organization of Z-discs (Zasp) and F-actin in more detail. *Dm*CFL-knockdown larvae exhibited a range of muscle phenotypes at the 3^rd^ instar stage, which we categorized into 3 classes based on the extent of structural disruption (Figures 2A and 2B). Class 1 muscles (65.875±3.76%) resembled control muscles, as both Zasp and F-actin retain their sarcomeric organization throughout the fiber. Class 2 muscles (6.633±0.074%) displayed dense accumulations of F-actin at the fiber poles, although Zasp maintained normal sarcomeric organization and did not co-localize with the actin aggregates. Class 3 muscles (27.492±3.69%) were the most severely affected and showed a complete disorganization of F-actin and Zasp. In addition, we found protein aggregates of varying sizes in Class 3 muscles, which contained F-actin as well as Zasp proteins (Figure 2C). This detailed analysis showed that, in our model, individual muscles were differentially affected in *Dm*CFL knockdown larvae; the most severe phenotype involved the formation of protein aggregates at different positions throughout the muscle fibers.

**Figure 2:**
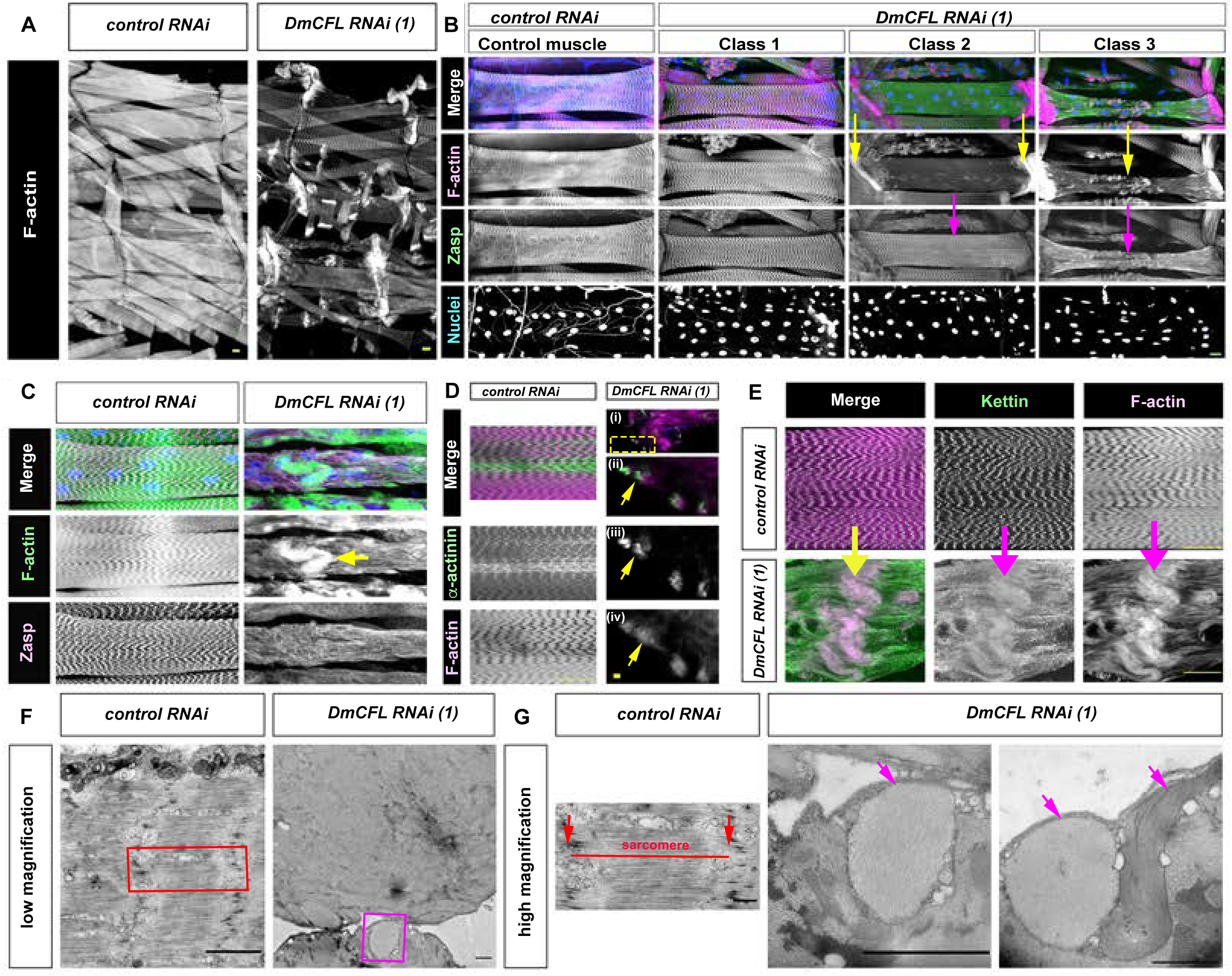
*Dm*CFL knockdown results in three classes of muscle phenotypes with varying protein aggregate composition. Larval muscles at the 3^rd^ instar stage expressing either *control RNAi* or *DmCFL RNAi (1)* driven with *Mhc-GAL4*. (A) Lateral view of a single larval hemisegment. Muscles are stained with Phalloidin (F-actin). *DmCFL*-knockdown larvae show a variety of muscle defects. (B) Classification of muscle phenotypes using VL3 muscles, labeled with Phalloidin (F-actin, magenta), α-Zasp (Zasp, green) and Hoechst (Nuclei, blue). Class 1 muscles appear similar to control muscles. In Class 2 muscles, F-actin is enriched at the muscle poles (yellow arrows), while Zasp retains its wild-type localization (magenta arrow). In Class 3 muscles, both actin and Zasp are disorganized (yellow and magenta arrows, respectively). Classification is based on N = 200 muscles from 8 larvae per genotype. (C-G) Protein aggregates in Class 3 muscles. (C-E) Class 3 muscle cells show severe sarcomere disorganization and the formation of protein aggregates containing F-actin (phalloidin) and the Z-disc proteins α-actinin (D, green) and Kettin (E, green). Arrows indicate protein aggregates. (C) Column 1: control muscle cell. Column 2: *DmCFL RNAi (1)* Class 3 muscle cell. Actin aggregates co-localize with Zasp. (D)Column 1: control muscle cell. Column 2: *DmCFL RNAi (1)* Class 3 muscle cell (i): Low magnification image of actin aggregates, yellow box denotes region magnified in subsequent panels (ii)-(iv). (ii)-(iv): Single Z slices taken from (i) panel. Actin aggregates co-localize with α-actinin (yellow arrows). (E) Top row: control muscle cell. Row 2: *DmCFL RNAi (1)* Class 3 muscle cell. Actin aggregates colocalize with Kettin. (F-G) TEMs of larval muscles at low (F) and high (G) magnification. Control muscles show organized sarcomeres (indicated in red), bounded by electron dense (dark) Z-disc proteins. In muscle cell expressing *DmCFL RNAi (1)* sarcomeric organization can be completely disrupted and aberrant protein aggregates found in the cell periphery (indicated in purple). Scale bars: 25 µm (A-E), 2 µm (F), 1µm (G).

Nemaline bodies, the diagnostic feature of NM, are protein aggregates containing Z-disc proteins such as α-Actinin, along with other sarcomeric proteins including actin, tropomyosin, and nebulin (Agrawal et al., 2007; 2004; Ilkovski et al., 2001; Sewry et al., 2019). Nemaline bodies either diffusely distribute in the cytoplasm, cluster under the cell membrane, localize near nuclei, or in lines within muscle fibers (Sewry et al., 2019). To determine if the aberrant aggregates observed in our *Dm*CFL-knockdown muscles were nemaline bodies, we examined and found co-localization of the F-actin aggregates and α-Actinin in our *Drosophila* model (Figure 2D). In addition, we observed co-localization of the F-actin structures with other components of the Z-disc, such as Kettin, in the *Dm*CFL-knockdown muscle fibers (Figure 2E). However, upon examination of the ultrastructure of larval muscles by Transmission Electron Microscopy (TEM), we observed severe disruption of organized sarcomeres (Figure 2F) and the presence of actin and Z-disc aggregates near the muscle periphery (Figures 2F and 2G), which did not resemble the electron dense, rod-like nemaline bodies seen in NM patients (Agrawal et al., 2007; Malfatti et al., 2014).

Altogether, these data indicated that muscle-specific knockdown of *Dm*CFL leads to sarcomeric degradation, and to the formation of protein aggregates that shared some similarities to nemaline bodies seen in NM patients.

### *Dm*CFL knockdown muscles show progressive deterioration

The structural changes in *Dm*CFL-knockdown muscles described thus far were observed at the end of larval development, in late 3^rd^ instar larvae. To examine the development of these phenotypes over time, we analyzed sarcomere organization each day over the five-day larval period (Figure 3A). At the end of embryonic development (Stage 17), when sarcomeres are initially established, we did not detect significant differences between *Dm*CFL-knockdown and control larvae. Compared to control, *Dm*CFL-knockdown larvae showed perturbations of the sarcomeres starting at the 2^nd^ instar stage, which then continued to deteriorate throughout the 3^rd^ instar stages. This loss in sarcomere organization was accompanied by a steady decline in muscle function at the 2^nd^ and 3^rd^ instar stages, as measured by average larval crawling velocity (Figure 3B). Together, these data suggested that, while functioning muscles are initially established, the deterioration of sarcomere structure and muscle function is progressive and occurs in parallel over several days of *Dm*CFL knockdown.

**Figure 3:**
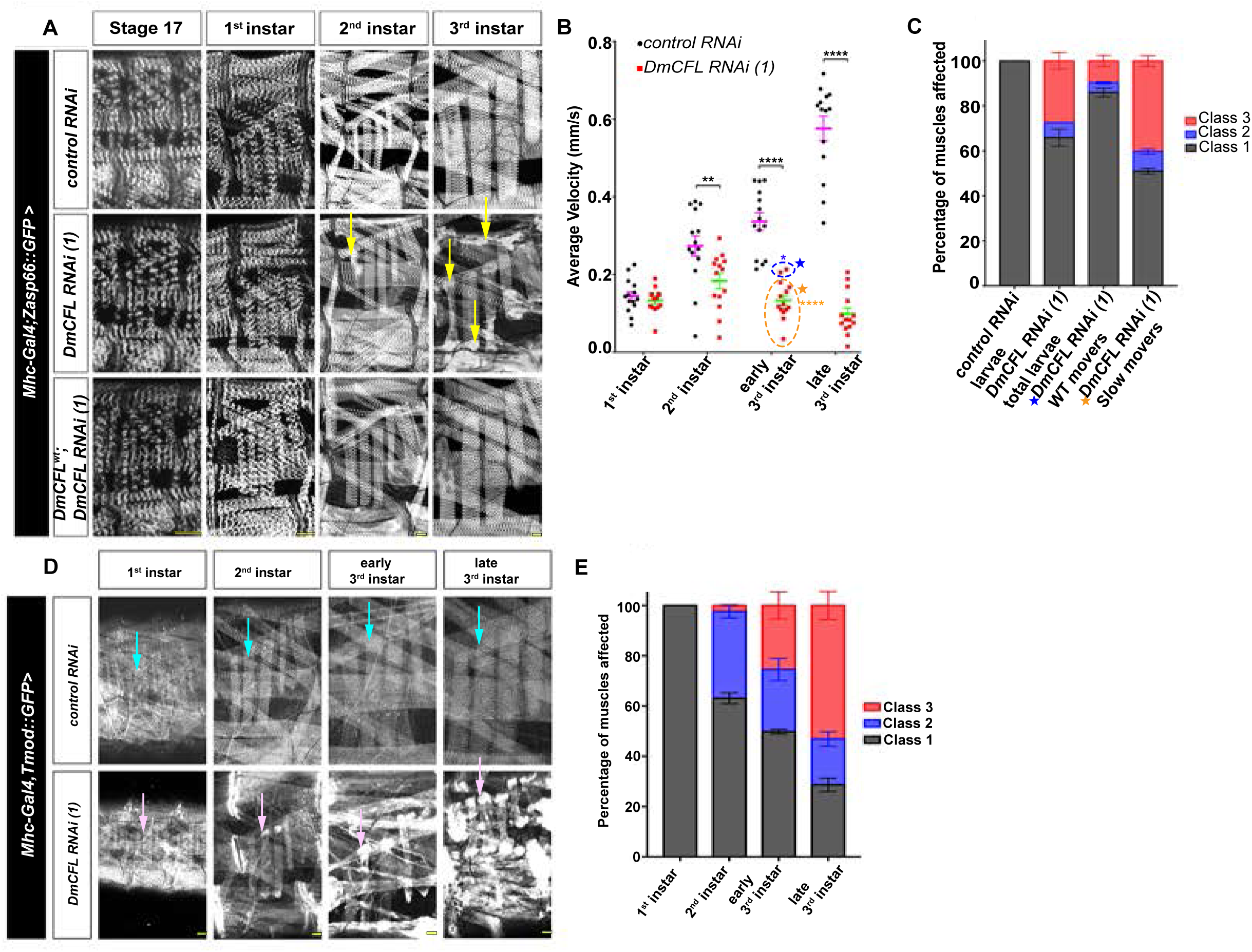
*Dm*CFL knockdown in the muscle results in progressive loss of both muscle structure and function. (A) Larval muscles over a 5-day period, from late embryogenesis (stage 17) through the different larval instars. Sarcomeres are labelled by Zasp-GFP (greyscale). Compared to *control RNAi*, *DmCFL RNAi* muscles show sarcomeric disorganization by 2^nd^ instar, and increased numbers of defects at the 3^rd^ instar stage (yellow arrows). Expression of *DmCFL^WT^* rescues this phenotype. (B) Crawling velocities of *control RNAi* and *DmCFL RNAi (1)* larvae at different larval stages (mean±SEM). Overall, *DmCFL*-knockdown significantly reduces larval speed (2-way ANOVA), beginning at the 2^nd^ instar stage. In early 3^rd^ instars, *DmCFL RNAi* larvae can be grouped into wild-type (WT) movers (blue dotted circle and star) and slow movers (orange dotted circle and star; Ordinary one-way ANOVA). *p<0.05, ***p<0.0002, ****p<0.0001. N = 15 larvae of each genotype repeated twice. (C) Percentage of different muscle classes seen in the specified genotypes at early 3^rd^ instar stage (mean±SEM). In comparison to *control RNAi* [100.0±0 %], *DmCFL RNAi (1)* muscles are represented by 3 classes [Class 1: 65.87±3.76%, Class 2: 6.63±0.07%, Class 3: 27.49±3.69%]. WT movers (B, blue star individuals) possess significantly more Class 1 muscles [Class 1: 85.94±1.95%, Class 2: 4.32±0.51%, Class 3: 9.72±2.46%], while slow movers (B, orange star individuals) show have more Class 3 muscles [Class 1: 50.95±1.21%, Class 2: 8.74±1.14%, Class 3: 40.29±2.36%]. N = 800 muscles, from at least 8 larvae of each genotype, and repeated twice. Statistical analysis in S2A. (D) Muscle cells from the same larva of each genotype imaged over 4 consecutive days. Sarcomeres are labelled with Tmod-GFP (greyscale). Individual *DmCFL RNAi (1)* muscles first develop into a Class 2 and ultimately a Class 3 muscle (magenta arrows highlight one example). (E) Percentage of muscle classes in *Dm*CFL depleted larvae at each stage of larval development (mean±SEM). 1^st^ instar [Class 1: 100±0.0%], 2^nd^ instar [Class 1: 63.11±2.18%, Class 2: 34.47±2.5%, Class 3: 2.42±0.32%], early 3^rd^ instar [Class 1:49.81±0.94, Class 2: 24.76±4.40%, Class 3: 25.42±5.34%] and late 3^rd^ instar [Class 1: 28.67±2.63%, Class 2: 18.20±2.93, Class 3: 53.121±5.56%]. N=200 muscles, from at least 5 larvae of each genotype, repeated twice. Statistical comparison in S2B. Scale bars: 25 µm (A, D).

To further probe the link between the deterioration of muscle structure and function, we analyzed the locomotive performance and the distribution of muscle classes in individual *Dm*CFL-knockdown larvae at the early 3^rd^ instar stage. We found that the larvae at the top of the functional range had a higher percentage of Class 1 and lower percentage of Class 3 muscles [Class 1: 85.94±1.95%, Class 2: 4.32±0.51%, and Class 3: 9.72±2.46%] compared to the larvae at the bottom of the functional range [Class 1: 50.95±1.21%, Class 2: 8.74±1.14%, and Class 3: 40.29±2.36%] (Figure 3C; S2A). This further supported a direct correlation between defective muscle structure and impaired muscle function.

To confirm that the muscle classes that we identified represent different stages in the progression of *Dm*CFL-dependent muscle deterioration, we imaged the musculature of individual larvae over the course of larval development. These analyses clearly showed that individual fibers that started out as Class 1 muscles progressed to Class 2 and culminated as Class 3 (Figure 3D). We did not detect *Dm*CFL protein in either Class 2 or Class 3 muscle cells by immunofluoresence, suggesting that this progression was not the result of a sudden depletion of *Dm*CFL at this stage (S2C). Quantification of the different muscle classes throughout larval development (Figure 3E, S2B) further established a progressive increase in the number of Class 2 and 3 muscles with successive larval stages.

Together, these data suggested that the progressive deterioration of *Dm*CFL-knockdown muscle fibers follows a specific pattern of structural changes of sarcomeric proteins that directly correlates with locomotive performance.

### F-actin aggregates at muscle poles recruit sarcomeric Tropomodulin and Troponin

We next sought to understand the temporal sequence in which the sarcomeric structure deteriorates upon *Dm*CFL knockdown. To this end, we examined the localization of other sarcomeric proteins in the different classes of *Dm*CFL-knockdown muscle fibers. All proteins tested were correctly localized in Class 1 muscle fibers and mislocalized in Class 3 muscles (S3A-S3G), yet showed distinct patterns in Class 2 muscles, which exhibit the first overt structural and functional changes. Proteins that maintained their localization in Class 2 muscle cells included the actin regulating protein Lasp/ Nebulin, the barbed end binding protein Cpa/CapZ, the actin polymerizing protein SALS, Myosin heavy chain (Mhc), and the M-line protein Obscurin (S3B and S3D-S3G). In contrast, Tropomodulin (Tmod) and Troponin T (TnT), two proteins with capping function at the pointed ends of actin filaments, showed aberrant patterns in Class 2 muscle cells (Figures 4A and 4B). In control muscle cells, Tmod localized in a repeated manner at the pointed end of the actin filaments, while TnT localized along the length of the sarcomeric actin filaments, as previously described (Farah and Reinach, 1995; Mardahl-Dumesnil and Fowler, 2001). In Class 2 muscle cells, both Tmod and TnT co-localized with F-actin at the cell poles and their levels were decreased in the center of the muscle cells (Figures 4C-4F). However, Tmod and TnT intensities within the entire Class 2 muscle fibers were not upregulated (S3H and S3I). These data indicated that in *Dm*CFL-knockdown muscle fibers Tmod and TnT were recruited from their association with existing sarcomeres at the center of the cell to the sites of actin accumulation at the cell poles.

**Figure 4:**
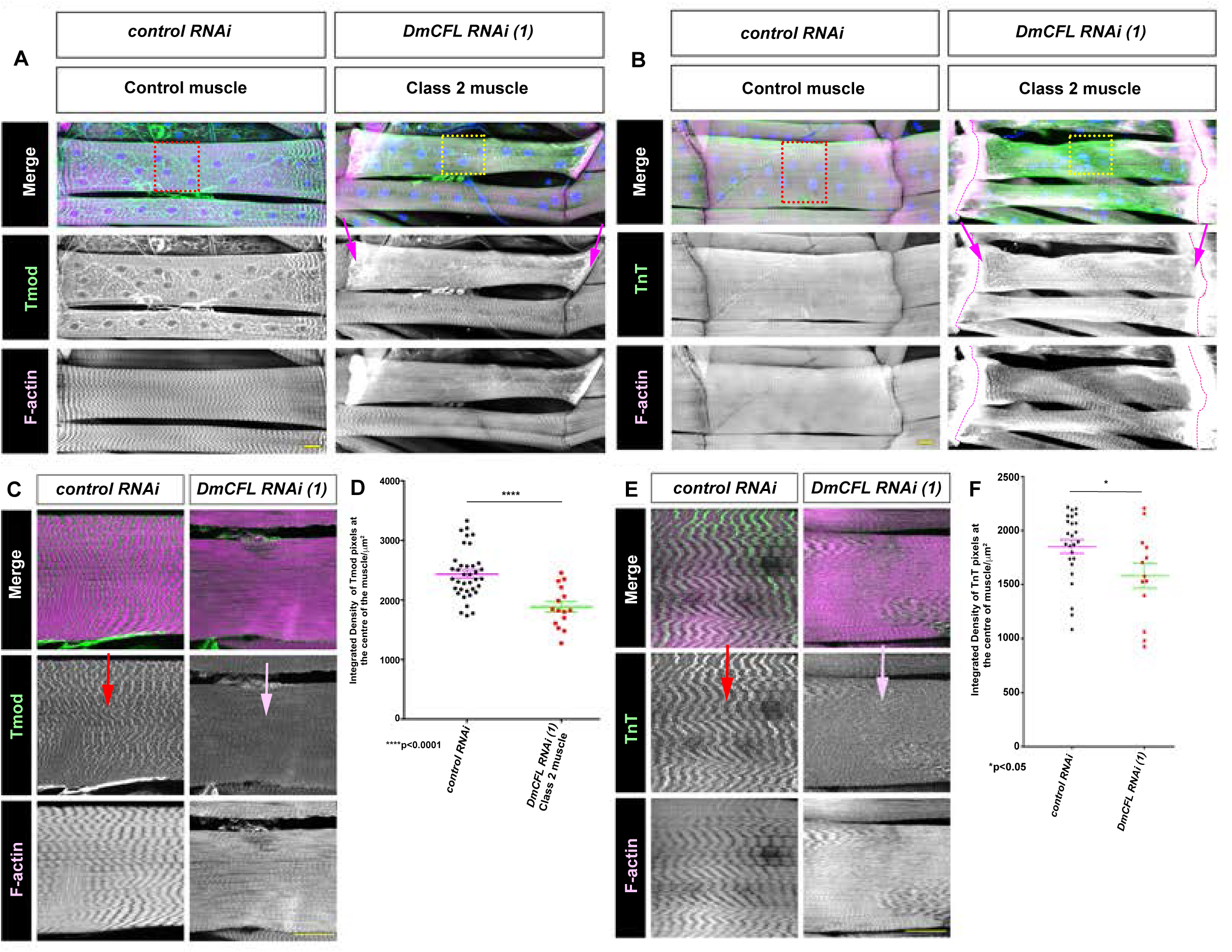
*Dm*CFL knockdown results in recruitment of sarcomeric Tropomodulin and Troponin to actin aggregates. Analysis of Tropomodulin (Tmod), Troponin (TnT) and Phalloidin (F-actin, magenta) protein localization in VL3 muscle from 3^rd^ instar larvae expressing either *control RNAi* or *DmCFL RNAi (1)* driven with *Mhc-GAL4*. (A) Tmod (α-Tmod) localizes to the sarcomeres throughout the control RNAi muscle cells. In *DmCFL*-knockdown Class 2 muscles, Tmod is found in sarcomeres but also co-localizes with actin at the muscle cell poles (magenta arrows). Dotted red and yellow boxes are the regions magnified in (C) and quantified (D). (B) TnT (α-TnT) localizes to the sarcomeres throughout the control RNAi muscles. In *DmCFL* knockdown Class 2 muscles, TnT is found in sarcomeres but also co-localizes with actin at the muscle cell poles (magenta arrows). Dotted pink lines demarcate muscle ends. Dotted red and yellow boxes are the regions magnified in (E) and quantified in (F). (C) Close up of muscle fibers shown in (A). In controls, Tmod localizes to the F-actin free region of the sarcomeres i.e. M-line (red arrows). In Class 2 *DmCFL*-knockdown muscles, sarcomeric Tmod levels are low (pink arrow; quantified in (D)). (D) Quantification of Tmod at the center of the muscle cells (pixel intensity values/µm^2^). Example images are shown in (C). P values calculated by Unpaired t-test (****p<0.0001). Error bars, mean ±SEM. N ³10 muscles from at least 5 different larvae, repeated twice. (E) Close up of muscle fibers shown in (B). In controls, TnT localizes at the pointed ends of the actin filaments in the sarcomeres (red arrow). In Class 2 *DmCFL*-knockdown muscles, TnT levels in the sarcomeres are lower (yellow arrow, quantified in (F)). (F) Quantification of TnT at the center of the muscle cells (pixel intensity values/µm^2^). Example images are shown in (E). P values calculated by Unpaired t-test (*p<0.05). Error bars, mean±SEM. N ≥10 muscles from at least 5 different larvae, repeated twice. Scale bars: 25 µm (A-C,E)

Together these data indicated that the ectopic accumulation of actin and the recruitment of pointed-end actin capping proteins represent the first visible intracellular signs of deterioration in our *Dm*CFL-knockdown model.

### *Dm*CFL-knockdown muscles form F-actin aggregates instead of new sarcomeres

The onset and progression of the phenotypes in *Dm*CFL-knockdown muscle cells coincide with a rapid growth phase of the musculature during the late 2^nd^ and early 3^rd^ instar stages (Figure 1). During this period, control muscles increased by 50% in both muscle cell length and sarcomere number, while a significant number of *Dm*CFL-knockdown muscles started accumulating F-actin at their poles (Figure 2). The poles of muscle fibers have long been thought to be sites of sarcomerogenesis in both skeletal muscle fibers (Bai et al., 2007; Dix and Eisenberg, 1990a; HAAS, 1950), as well as cardiomyocytes (Yang et al., 2016). To determine whether *Dm*CFL knockdown affects the addition of new sarcomeres during muscle cell growth, we specifically analyzed pairs of VL muscles in which one cell had a Class 1 (control-like) phenotype while the other was Class 2 (Figures 5A-5B). We found that, despite having the same length, Class 2 muscles contained significantly fewer sarcomeres compared to their Class 1 neighbors (Figures 5C and 5D; S4A).

**Figure 5:**
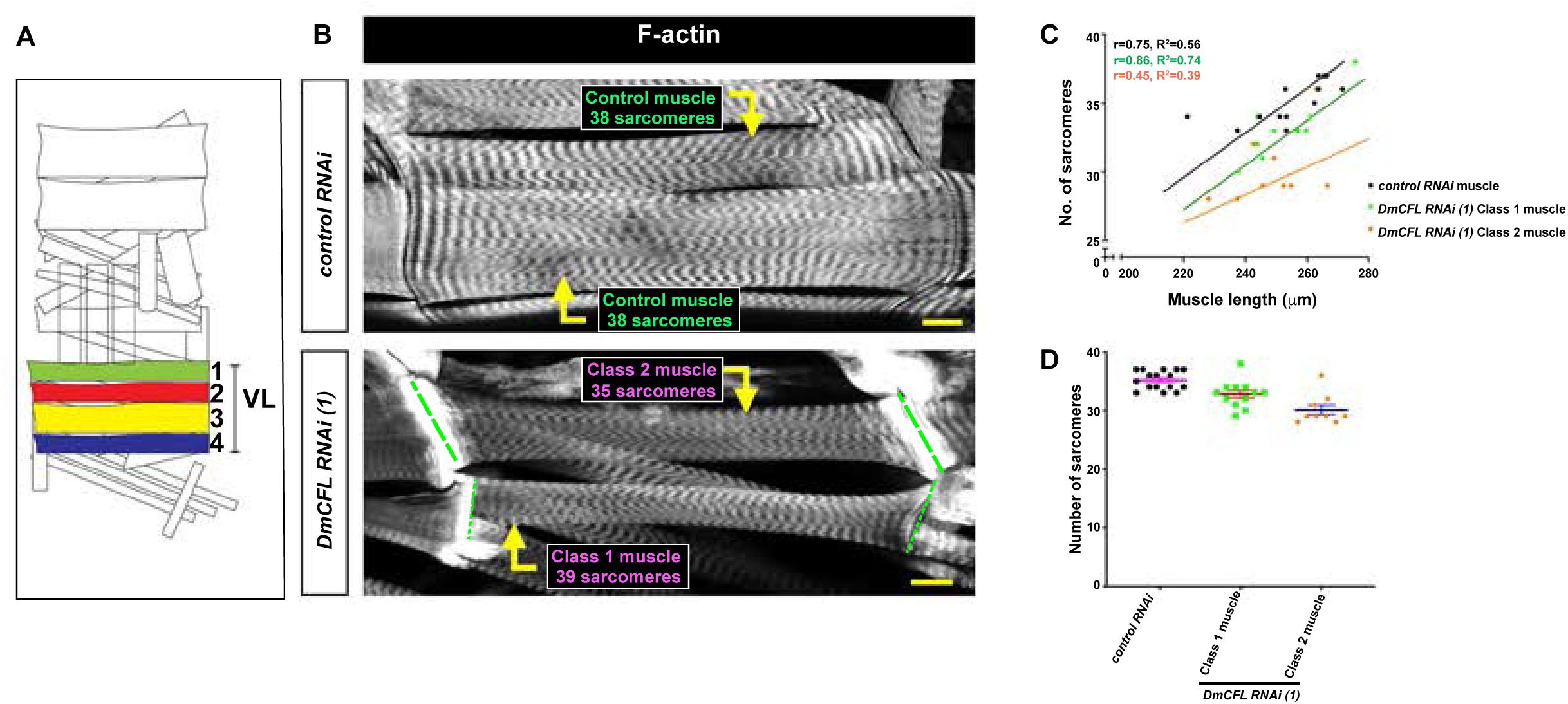
*Dm*CFL knockdown leads to fewer sarcomeres being added in growing muscle ends. (A) Diagram of one hemisegment of the larval musculature, with the VL muscles highlighted in different colors. (B) VL muscles from larvae expressing either *control RNAi* or *DmCFL RNAi (1)* and stained with Phalloidin for F-actin (greyscale). In controls, different VL muscles of the same length have 38 sarcomeres each. In *Dm*CFL knockdown muscles of the same length, a Class 1 VL muscle contains 39 sarcomeres, while a Class 2 VL muscle contains only 35 sarcomeres. Dotted lines demarcate muscle ends. Scale bar, 25μm (C) Quantification of muscle cell length (μm) and number of sarcomeres in VL muscles from *control RNAi* and *DmCFL RNAi (1)* expressing 3^rd^ instar larvae. Pearson’s correlation coefficient (r) and R^2^ values and best fit lines were calculated for each muscle cell class. (D) Number of sarcomeres in VL muscles shown in Figure 4C of the indicated genotypes. Error bars, mean±SEM. N ≥12 muscles from at least 5 different larvae. Statistical analysis in Supplementary Figure 4A.

These data suggested that *Dm*CFL plays a crucial role in the addition of new sarcomeres in growing muscle cells. In *Dm*CFL-knockdown muscles, F-actin forms aggregates at the cell poles instead of contributing to the formation of new sarcomeres.

### Increased proteasome activity improves *Dm*CFL knockdown phenotypes

*Dm*CFL knockdown in the muscle fibers resulted in the presence of large sarcomeric protein aggregates and overall deterioration of muscle structure and function. We next asked why sarcomeric proteins accumulated instead of being degraded by the proteolytic machinery (Figure 6A). We did not observe a significant difference in overall Zasp-GFP recovery rates in either situation (Figure 6B). Furthermore, the levels of the mobile fractions of Zasp-GFP were similar in both *control RNAi* and *DmCFL RNAi (1)* expressing muscle cells (Figure 6C). These data suggested that the sarcomeric protein aggregates in the *Dm*CFL-knockdown muscle fibers, like the sarcomeres in control fibers, are dynamic structures. Furthermore, the levels of the mobile fractions of Zasp-GFP were similar in both *control RNAi* and *DmCFL RNAi (1)* expressing muscle cells (Figure 6C). Together, these data indicated that the sarcomeric protein aggregates in the *Dm*CFL-knockdown muscle fibers, like the sarcomeres in control fibers, are dynamic structures with proteins rapidly binding and being released.

**Figure 6:**
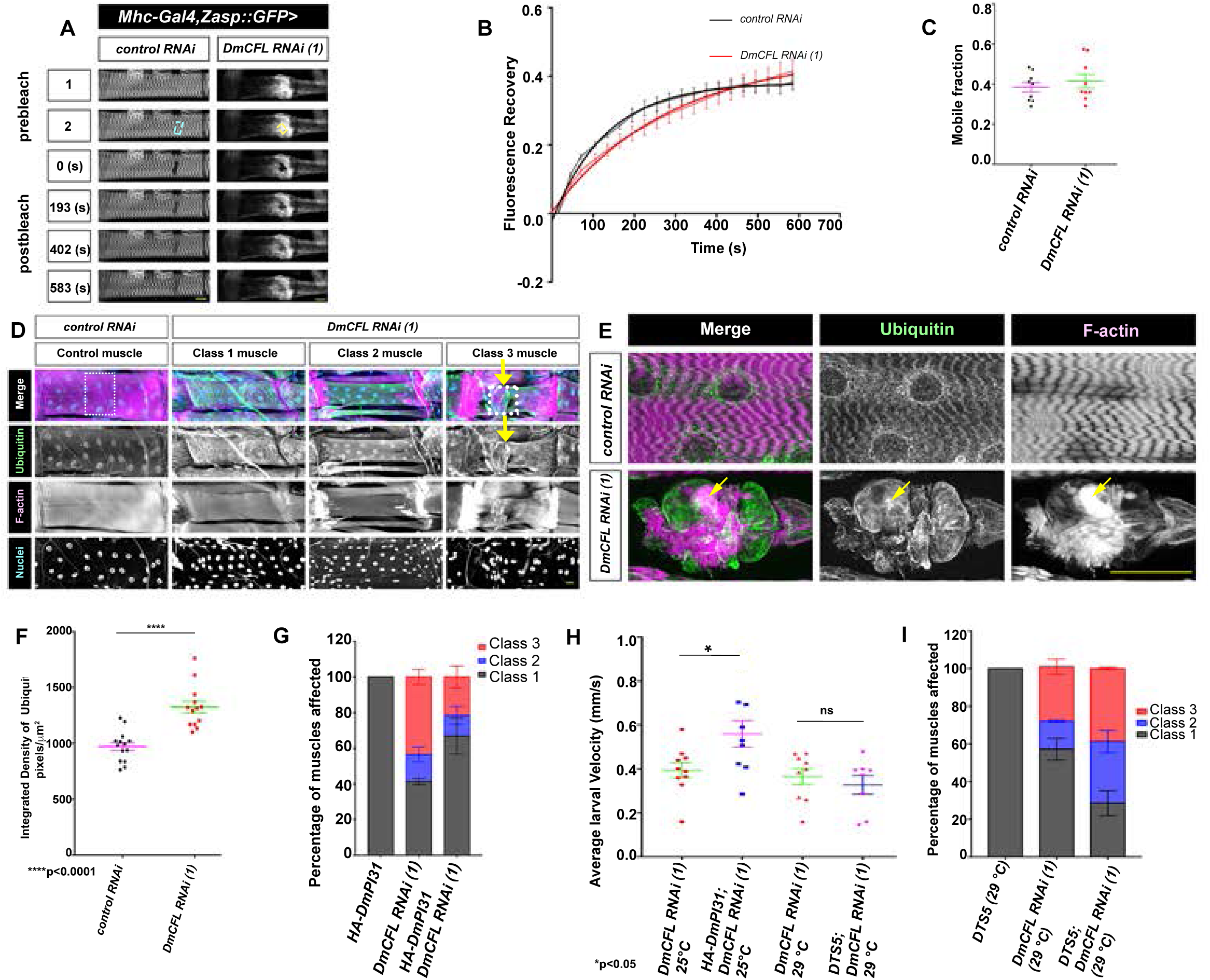
Modulating proteasome activity affects the progression of structural and functional changes in *Dm*CFL-knockdown muscles. (A) Single muscles expressing Zasp-GFP(greyscale) with either *control RNAi* or *DmCFL RNAi(1)* before and after Fluorescence Recovery After Photobleaching (FRAP). Dotted rectangles outline photobleached region. No significant difference in overall recovery rate. (B) Quantification of Zasp-GFP FRAP in *control RNAi* or *DmCFL RNAi (1)* expressing 3^rd^ instar muscle cells. N=9 muscles per genotype, repeated twice. mean ±SEM (C) Mobile fraction analysis of Zasp-GFP present in *control RNAi* and *DmCFL RNAi (1)* larvae. P values calculated by unpaired t-test (p=0.685, not significant). (D) VL3 muscles from 3^rd^ instar larvae expressing either control RNAi or *DmCFL RNAi (1)*. Labeling for the presence of mono-and poly-ubiquitin (α-FK2, green), F-actin (Phalloidin, magenta), and nuclei (Hoechst, blue). In the Class 1 and 2 muscle cells (second and third columns, respectively), FK2 localizes in the same pattern seen in the *control RNAi* muscle cells. In the Class 3 muscle cells (fourth column), FK2 levels are increased (yellow arrows). N=8 larvae of each genotype repeated in triplicate. Dotted boxes are areas magnified in (E). (E) High magnification of muscles as shown in (D). Ubiquitin in *control RNAi* muscles localizes around the nuclei and in the Z-discs. In *DmCFL RNAi (1)* Class 3 muscles, FK2 is expressed at higher levels (quantified in (F)) and co-localizes with sarcomeric protein aggregates (yellow arrows). (F) Quantification FK2 levels (pixel intensity values/µm^2^) in *control RNAi* and Class 3 *DmCFL RNAi (1)*-expressing muscles (mean± SEM). P values calculated by unpaired t-test (**** p < 0.0001). N ≥13 muscles from at least 5 different larvae, repeated twice. (G) Percentage of different muscle classes in the specified genotypes. In comparison to *DmCFL RNAi (1)* [Class 1:41.403 ±1.68%, Class 2:15.09 ± 4.17%, Class 3 :43.5 ±4.11%], expression of *HA::DmPI31* in *Dm*CFL-depleted muscle cells increases the number of Class 1 and 2 muscle cells and decreases the number of Class 3 muscles [Class 1:66.87±10.05%, Class 2: 11.59 ± 5.08%, Class 3: 21.52 ± 6.12%]. Error bars, mean ±SEM. N = 800 muscles, from at least 8 larvae of each genotype, repeated twice. Statistical analysis shown in Supplementary Figure 5E. (H) Crawling velocities of 3^rd^ instar larvae of the specified genotypes. P values by Unpaired *t*-test (* p<0.05, ns p=0.498). Expression of *HA::DmPI31* with *DmCFL RNAi (1)* significantly improves velocity; expression of *DTS5* with *DmCFL RNAi (1)* does not significantly affect velocity. Error bars mean± SEM. N=9 larvae per genotype, repeated twice. (I) Percentage of different muscle classes in the specified genotypes. In comparison to *DmCFL RNAi (1)* [Class 1: 57.14 ± 5.7%, Class 2: 14.8 ± 0.68%, Class 3: 28.99 ± 4.09%] expression of *DTS5* with *DmCFL RNAi,* decreases the number of Class 1 muscles and increases the number of Class 2 and Class 3 muscles [Class 1: 28.53 ± 6.62 %, Class 2: 32.72 ± 5.9%, Class 3: 38.74 ± 0.72%]. Expression of *DTS5* in a *control* background does not impact muscle structure (see also Supplementary Figures 5F and 5H). Error bars, mean ±SEM. N = 800 muscles, from at least 8 larvae of each genotype, repeated twice. Statistical comparison of each muscle class for each genotype shown in Supplementary Figure 5E. Scale bars: 25 µm (A,D,E)

To investigate the possibility that the sarcomeric protein aggregates found in *Dm*CFL-knockdown muscle cells result from defective protein degradation, we manipulated protein degradation by the proteasome. As the proteasome targets and degrades ubiquitinated proteins, we first examined whether mono-and poly-ubiquitination of the protein aggregates occurred. We found that the sarcomeric proteins in the aggregates were indeed ubiquitinated (Figures 6D-6E). Further, overall ubiquitin levels were elevated in the Class 3 muscle cells compared to controls (Figure 6F), indicating that the aggregated proteins were properly targeted for degradation.

To test the ability of the muscle cells to degrade the ubiquitinated proteins, we overexpressed DmPI31, which increases the activity of the proteasome (Bader M et al., 2011). In control muscles, DmPI31 overexpression had no obvious effect on muscle structure, function, or viability of the organism (S5A-S5D). Immunofluorescence images of the larval muscles expressing *DmPI31* with *DmCFL RNAi (1)* indicated that DmPI31’s localization in both Class 1 and 2 muscle cells was similar to that seen in the control muscle cells (S5B). However, co-expression of *DmPI31* with *DmCFL RNAi (1)* resulted in a significant improvement in muscle cell structure and function compared to expression of *DmCFL RNAi (1)* alone (Figures 6G and 6H, S5E). Most striking, the percentage of Class 1 muscle cells was increased at the cost of Class 3 muscles, indicating that the progression of sarcomeric disassembly was delayed by increasing proteasome activity.

To confirm this result, we inhibited proteasomal activity by expressing *DTS5*, a temperature sensitive missense allele of the *β*6 subunit of the 20S proteasome, which inhibits proteasomal activity by acting as a dominant negative (Belote and Fortier, 2002) (Belote JM and Fortier E, 2002). Embryos expressing these constructs were moved to the restrictive temperature (29°C) and remained there throughout larval development. We observed no change in muscle cell structure and function upon DTS5 expression in the control muscles (S5C, S5F, and S5G). However, *DTS5* expression combined with *DmCFL RNAi (1)* caused accelerated muscle cell degeneration, with an increase in aberrant muscle structure (Figure 6I, Class 3: 38.74 ± 0.72%). We did not detect any further worsening in larval locomotion velocity, which was already at a very low level. Together these data indicated that muscle cell degeneration was accelerated by inhibiting the proteasome (Figure 6H). Expression of *DmPI31* or *DTS5* in *DmCFL-*knockdown muscle cells did not affect the depletion of *Dm*CFL by *DmCFL RNAi (1)* and failed to improve the overall viability (S5G and S5I).

Together, these results suggested that in our *Dm*CFL-knockdown model, manipulation of the proteasome and can delay the progression of structural and functional muscle deterioration and the formation of protein aggregates.

### NM Patient point mutation in CFL2 abrogates CFL function *in vitro* and *in vivo*

*CFL2* mutations seen in NM patients include both small deletions and point mutations(Agrawal et al., 2007; Ockeloen et al., 2012; Ong et al., 2014). Our *Dm*CFL-knockdown models recapitulates some aspects of muscle cell degeneration observed in a severe case of human NM, in which a *CFL2* null mutation was detected (Ong et al., 2014). This patient’s muscle fibers showed nemaline bodies at the cell peripheries and reduced muscle function. Point mutations in *CFL2,* thought to be hypomorphs, have been identified in patients with less severe cases of NM. One of these point mutations alters a single amino acid p.Val7Met(V7M) that is conserved between *Drosophila*, mouse, and human Cofilins/Tsr (Figure 7A, S6A). One of these point mutations alters a single amino acid p.Val7Met(V7M) that is conserved between *Drosophila*, mouse, and human Cofilins/Tsr (Figure 7A, S6A). To understand how the V7M mutation affects protein function, we biochemically compared the interactions and activities of human CFL2^WT^ and CFL2^V7M^. Analysis of the nucleotide exchange rate on actin monomers (ADP to ATP) revealed that the rate is increased in CFL2^V7M^, suggesting that it has enhanced affinity for G-actin (Figure 7B). Further, *in vitro* TIRF microscopy analysis revealed that the ability of CFL2^V7M^ to sever actin filaments is compromised, evident from its reduced plateau of cumulative severing compared to CFL2^WT^ (Figures 7C and 7D, S6B). Continuous actin turnover is critical for maintaining sarcomere structure and muscle function (S. Ono, 2010). Our experiments showed that the V7M mutation in CFL2 severely compromises CFL2’s functional interactions, by increasing the levels of ATP-actin monomers and decreasing F-actin severing rate.

**Figure 7:**
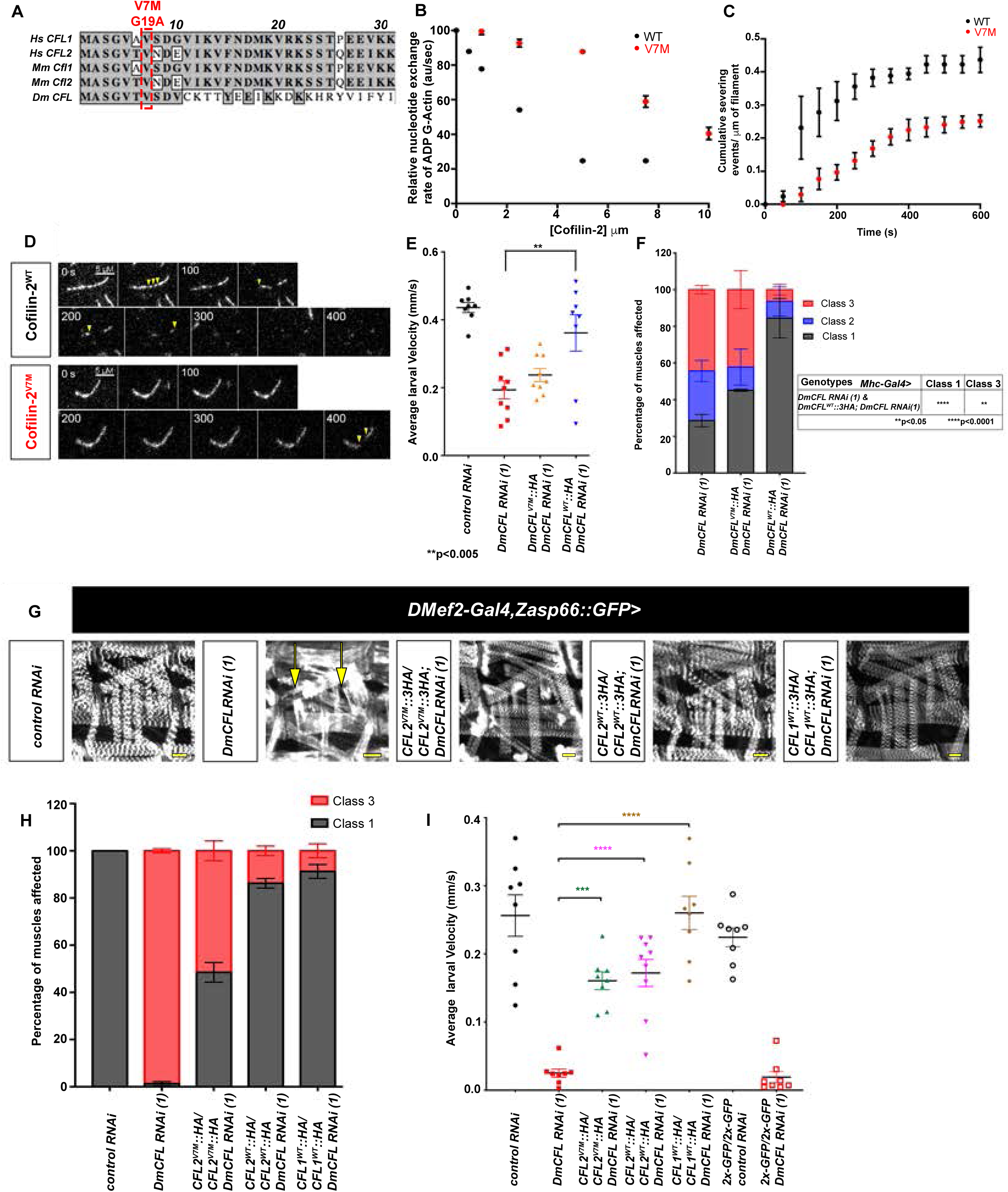
*In vivo* and *in vitro* analyses identify phenotypic and biochemical consequences of a specific patient *CFL2* mutation. (A) Sequence alignment of proteins encoded by human Cofilin-1 (*HsCFL1*) and Cofilin-2 (*HsCFL2*), mouse *MmCfl1* and *MmCfl2*, and *Drosophila DmCFL* genes. Sequence alignment of the first 30 amino acids indicates strong conservation (grey; for entire protein sequences see Supplementary Figure 6A). Val7 is a conserved amino acid in all 5 proteins. Val7 in *Hs*CFL2 was found mutated to Met in patients with NM, Val7Met (V7M). (B) Comparison of nucleotide exchange rate on 2 μM ADP-actin monomers in the presence of different concentrations of human CFL2^WT^ and CFL2^V7M^. Error bars, mean±SEM. (C) Analysis of actin filament severing activity. Each data point is the number of cumulative severing events per micron of F-actin at that time point, averaged from three independent experiments (N = 20 actin filaments each). Error bars, mean±SEM. (D) Representative time-lapse images from TIRF microscopy experiments in which tethered OG-labeled actin filaments were preassembled, and then 150 nM CFL2 ^WT^ or CFL2^V7M^ was flowed in. Yellow arrowheads indicate severing events. (E) Crawling velocities of late 3^rd^ instar larvae with the indicated genotypes. P values were calculated by Ordinary one-way ANOVA (**p<0.005). *DmCFL^WT^* rescues larval locomotion in comparison, while *DmCFL^V7M^* fails to significantly rescue this phenotype. N=9 larvae of each genotype, repeated at least twice. **p<0.005. Error bars, mean±SEM. (F) Percentage of different muscle cell classes in the specified genotypes. In comparison to *DmCFL RNAi (1)* knockdown individuals [Class 1: 28.57 ± 3.38%, Class 2: 27.10 ± 5.75%, Class 3: 44.31 ± 2.37%], co-expression of *DmCFL^WT^* with *DmCFL RNAi* increases the number of Class 1 muscle cells and decreases the number of Class 3 muscles [Class 1: 84.44 ± 10.73%, Class 2: 9.13 ± 7.90%, Class 3: 6.42 ± 2.83%]. Coexpression of *DmCFL^V7M^* with *DmCFL RNAi (1)* partially rescues *DmCFL* depletion [Class 1: 45.11 ± 0.57%, Class 2: 12.66 ± 9.83%, Class 3:42.21 ± 10.41%]. N= 800 muscles, from at least 8 larvae of each genotype, repeated twice. P values were calculated by two-way ANOVA (**p<0.05, ****p<0.0001). Error bars, mean±SEM. (G) Muscles of 2^nd^ instar larvae of the specified genotypes. Sarcomeres are labeled by Zasp-GFP expression (greyscale). In *DmCFL RNAi* muscles, sarcomeres are highly disorganized (yellow arrows). Expression of high levels (two copies) of *HsCFL2^V7M^* improves the muscle phenotype. Expression of high levels (two copies) of *HsCFL2^WT^* or *HsCFL1^WT^*, almost completely rescues sarcomere defects. Bar, 25um. (H) Percentage of different muscle cell classes in the specified genotypes. In comparison to *DmCFL RNAi (1)* alone [Class 1: 1.294 ± 0.887%, Class 3: 98.706 ± 0.887%], co-expression of *HsCFL2^WT^* significantly rescues muscle defects [Class 1: 86.249 ± 2.025, Class 3:13.751 ± 2.025%]. Similar levels of rescue are also seen with *HsCFL1^WT^* [Class 1: 91.251 ± 2.930, Class 3:8.749± 2.930%]. Co-expression of *HsCFL2^V7M^* shows a slight improvement [Class 1: 48.503 ± 4.185%, Class 3: 51.497 ± 4.185%], consistent with this hypomorphic mutation. N= 600 muscles, from at least 8 larvae of each genotype, repeated twice. Statistical analysis shown in Supplementary Figure 6J. Error bars mean ±SEM. (I) Crawling velocities of 2^nd^ instar larvae in which *Dmef2-Gal4* is used to express the constructs as listed. Muscle-specific expression of *HsCFL2^V7M^, HsCFL2^WT^, and HsCFL1^WT^* with *DmCFL RNAi (1*) significantly rescues larval locomotion compared to *DmCFL RNAi (1)* larvae alone. Co-expression of two copies of 2x-GFP with *DmCFL RNA (1)* does not rescue larval velocity. There was no significant difference in the velocities between *control RNAi* and *2x-GFP/2x-GFP; control RNAi* larvae (ns., p=0.355) or between *DmCFL RNAi (1)* and *2x-GFP/2x-GFP; DmCFL RNAi (1)* larvae (ns., p=0.569). N=8 larvae of each genotype, repeated twice. P values were calculated by Ordinary one-way ANOVA (***p<0.0002, ****p<0.0001). Error bars, mean ±SEM. Statistical analysis shown in Supplementary Figure 6J.

To test the effects of the V7M mutation in *Drosophila* muscles, we engineered UAS expression constructs for both *DmCFL* and human *CFLs* containing the mutation (Figure 7A, S6A). Both *Dm*CFL^WT^ or *Dm*CFL^V7M^ localize to the Z-discs and H zones of the sarcomere, similar to endogenous *Dm*CFL in control muscle fibers (S6C-S6F, compare Figure 1G). While the expression of *DmCFL^WT^* in *DmCFL*-knockdown muscles rescued muscle function, muscle structure, and viability, expression of *DmCFL^V7M^* did not rescue wildtype *Dm*CFL function (Figures 7E and 7F, S6G). The expression levels of both constructs were confirmed through western analysis (S6H and S6I), suggesting that, like human CFL2^V7M^, *Dm*CFL^V7M^ is a hypomorph.

To test the function of human CFL proteins in our *Drosophila* model, we expressed *CFL2^WT^*, *CFL2^V7M^*, and *CFL1^WT^* in *Dm*CFL-knockdown larval muscle cells. We used the strong muscle GAL4 driver, *Dmef2-Gal4* and expressed two copies of each of the constructs. We confirmed expression levels of all constructs through western blots (S6J) and found that the proteins encoded by *CFL2^V7M^, CFL2^WT^*, and *CFL1^WT^* localized to the Z-disc and H-zone of the *Drosophila* sarcomere, similar to the fly protein (S6K-S6P). Co-expression with *DmCFL RNAi(1)* showed that *CFL2*^WT^ improved muscle cell structure and function (Figures 7G-7I), indicating that human CFL2^WT^ can rescue *Dm*CFL-knockdown in *Drosophila.* Expression of *CFL2^V7M^* in a similar manner also rescued both muscle cell structure and function, but not to the same extent as seen with CFL2^WT^ (Figures 7G-7I). We hypothesized that this rescue of muscle cell function and structure by CFL2^V7M^ can be attributed to the mutant protein retaining some function *in vivo* as was seen our *in vitro* experiments (Figures 7B-7D). Expression of *CFL1^WT^* in the *Dm*CFL-knockdown muscle cells also rescued muscle cell structure and function (Figures. 7G-7I). The slightly better rescue of CFL1 ^WT^, compared to that of CFL2^WT^, can be attributed to the higher level of expression (S6J). Expression of two copies of *UAS-GFP* along with *DmCFL RNAi (1)* did not improve larval muscle cell function (Figure 7I), indicating that Gal4 dilution is not contributing to improved muscle function seen in our rescue with *Dm*CFL and the human CFLs.

Together, these data showed close functional similarities between human and fly CFL proteins and indicated that the human V7M mutation impairs the ability of CFL2 to sever actin filaments both *in vivo* and *in vitro*.

## DISCUSSION

Sarcomeres are the fundamental force-producing, contractile units of skeletal muscles. While sarcomere structure and its composition have been studied extensively, how sarcomeres are assembled and added to a growing muscle remains unknown. Moreover, how mutations in sarcomeric proteins contribute to diseases, such as nemaline myopathy, is unclear. In this study, we show that *Dm*CFL, the *Drosophila* homolog of human CFL2, is required for both the addition and the maintenance of sarcomeres in muscle fibers of the *Drosophila* larva. We find that muscle-specific knockdown of *Dm*CFL results in progressive loss of sarcomere and muscle structure leading to loss of muscle function. *Dm*CFL knockdown further results in the formation of protein aggregates composed of actin and Z-disc proteins. We also show that progression of muscle structural and functional changes can be delayed by activation of the proteasome. Conversely, inhibition of the proteasome in *Dm*CFL knockdown muscles exacerbates the changes in muscle structure. Finally, we demonstrate close functional similarities between *Drosophila* and human Cofilin proteins both *in vitro* and *in vivo* and link a specific patient mutation in CFL2 to muscle phenotypes observed in NM patients.

Sarcomerogenesis and sarcomere homeostasis during muscle development have long been elusive processes. Both *in vivo* and *in vitro* studies have suggested that CFL2 localizes to the sarcomeres and is required for maintaining F-actin length and sarcomere structure during muscle development (Gurniak et al., 2014a; Kremneva et al., 2014b; K. Ono et al., 2003), which has been attributed to the actin severing function of CFL2 proteins (Andrianantoandro and Pollard, 2006; Michelot et al., 2007). We show that *Dm*CFL specifically localizes to both the Z-disc and H-zone in larval sarcomeres, and that muscle-specific knockdown of *Dm*CFL affects sarcomeric F-actin filaments. In the severely affected *Dm*CFL-knockdown muscles, we find a complete collapse of F-actin and sarcomere organization, and the formation of actin aggregates. The presence of F-actin accumulations in *Dm*CFL-knockdown muscles is consistent with previous studies where loss of CFL2 or its homologs results in F-actin rods/aggregates (Agrawal et al., 2012; Gurniak et al., 2014b; K. Ono et al., 2003; S. Ono et al., 1999; Subramanian et al., 2015), and in accordance with the role of CFLs in actin severing. Altogether, this study extends our understanding of CFL’s role sarcomere homeostasis and presents a *Drosophila* model highly suited for future studies investigating CLF function in the sarcomeres.

In addition to sarcomere homeostasis, our work uncovers the activity of CFL in sarcomerogenesis during muscle growth, an area which has not been studied previously in other model systems. The larval muscle grows 40-50-fold in size over a period of 5 days (Demontis and Perrimon, 2009), making this an outstanding system to study myofiber growth. We find that in controls, sarcomere numbers increase at each larval instar, correlating with an increase in muscle cell length. Loss of *Dm*CFL disrupts sarcomere addition in the growing muscles, resulting in F-actin accumulations at the poles coupled with fewer sarcomeres. This is particularly evident during the 2^nd^ to 3^rd^ instar transition, when control muscles experience the greatest rate of growth. TEM and live imaging data suggest that the ends of the myofibers are the sites of sarcomerogenesis (Dix and Eisenberg, 1990b; Yang et al., 2016). Further, the accumulation of F-actin at muscle cell poles has been previously observed in zebrafish models of NM (Sztal et al., 2015). Our data suggest that loss of *Dm*CFL leads to the accumulation of F-actin specifically in regions of sarcomere addition, thereby indicating a crucial role for *Dm*CFL in sarcomerogenesis.

Despite the critical role of *Dm*CFL in sarcomerogenesis, our knockdown model clearly showed that the deterioration of muscle structure and function were progressive in nature and occurred in parallel. Initially all *Dm*CFL-knockdown muscles had control features (Class1). These conditions are similar to *Cfl2* knockout mouse models, where, CFL2 is not necessary for initial myofibrillogenesis, but required for subsequent muscle growth and development (Agrawal et al., 2012; Gurniak et al., 2014b). On a cellular level, the decline in muscle function in the *Dm*CFL-knockdown larvae can be attributed to several factors. Firstly, fewer sarcomeres are added to the growing muscle, resulting in a weaker muscle with lower force production. Secondly, the formation of actin aggregates at the cell poles (Class 2 muscles), most likely a result of uncontrolled actin polymerization, could interfere with muscle contraction. Thirdly, the F-actin aggregates recruit Tmod and TnT, actin capping proteins, from existing sarcomeres. This mislocalization of Tmod is in accordance with a *Drosophila* model overexpressing *ACTA1* mutations seen in NM patients (Sevdali et al., 2013). The loss of Tmod and TnT from sarcomeres have been shown to result in reduced muscle function (Nworu et al., 2015; Singh et al., 2014; Stevenson et al., 2007). Also, in the absence of CFL2, capping proteins bind and regulate actin filament length less efficiently (Fowler, 1996; Kremneva et al., 2014a). It is likely that the suboptimal capacity of these capping proteins in combination with loss of *Dm*CFL contributes ultimately leads to the collapse of sarcomeric organization (Class 3 muscles), thereby significantly impairing muscle contraction and locomotion. Whether this progression is enhanced with increased muscle activity is an area for future study.

Sarcomeric proteins undergo rapid turnover during muscle contraction and myofibrillogenesis (Littlefield and Fowler, 2008; J. M. Sanger and J. W. Sanger, 2008; this study). We analyzed why aggregated proteins were not degraded by the proteasome and found that the aggregates were in constant flux and properly targeted for ubiquitination. Further we showed that upon proteasome overactivation, muscle deterioration was slowed down but not completely eliminated. These data suggested that the proteasome was unable to compensate for the lack of *Dm*CFL’s F-actin severing activity. While other severing proteins such as Gelsolins are expressed in the *Drosophila* muscle, they are unable to compensate for the loss of *Dm*CFL. Together these data further highlight the importance of *Dm*CFL for muscle structure and function. In addition, they provide crucial insights to the mechanisms underlying the formation of sarcomeric protein aggregates, as well as possible therapeutic approaches. The breakdown of sarcomeric protein aggregates and recycling of limiting components such as actin could improve muscle contraction and locomotion.

Mutations in CFL2 have been identified in three isolated cases of NM (Agrawal et al., 2007; Ockeloen et al., 2012; Ong et al., 2014). NM is a slowly progressing myopathy characterized by muscle weakness and the presence of nemaline bodies containing both actin and *α*-actinin (Sewry et al., 2019). Mouse models of *Cfl2* recapitulate aspects of the disease such as the degeneration of muscle fibers, presence of actin and *α*-actinin positive nemaline bodies, sarcomeric disruptions, actin accumulations and muscle weakness (Agrawal et al., 2012; Gurniak et al., 2014b). In our *Dm*CFL-knockdown *Drosophila* muscles, we see disruption of both muscle and sarcomere structure and loss of muscle function. These phenotypes can be rescued with the human CFL genes, indicating a strong conservation between the *Drosophila* and human proteins. While the protein aggregates in our *Dm*CFL-knockdown muscle fibers do not completely mimic the electron dense nemaline bodies seen in NM patients, they do have similar protein compositions (F-actin, Zasp, and *α*-actinin) and subsarcolemmal localization (Sewry et al., 2019).

Similar to NM patients and *Cfl2* mouse models (Agrawal et al., 2012; Gurniak et al., 2014b) not all muscle fibers in our *Dm*CFL-knockdown larvae and deteriorate at the same time and rate. Our data show that the timing and strength of RNAi expression correlate with the timing and severity of structural changes, indicating that protein levels play a crucial role in onset and progression of phenotypes. In our *Dm*CFL knockdown larvae, the progression of structural changes in the muscle fibers coincides with a period of significant muscle growth and sarcomerogenesis. Similarly, symptoms of NM have been known to develop during the prepubertal growth spurt (Sanoudou and Beggs, 2001). Mammalian muscle similarly grows in size postnatally and during puberty (Bachman et al., 2018; Lexell et al., 1992; Pearson, 1990).The postnatal period of muscle growth correlates with the transition from the expression of *CFL1* and *ADF* to *CFL2* genes (Abe and Obinata, 1989; Abe et al., 1989; Bamburg and Bray, 1987; Gurniak et al., 2014b). The expression of CFL2 as the predominant isoform in the skeletal muscle (Thirion et al., 2001) and the demand for increased sarcomerogenesis suggest that muscle growth could contribute to NM disease onset and progression. Using our *Drosophila* model to test whether factors like growth and muscle activity as modulators of muscle phenotypes remains a future goal.

In summary, our data strongly implicate a specific role for CFLs in sarcomere formation and maintenance through the regulation of actin thin filaments. Our work also suggests a model for nemaline body formation that rests on the fundamental structure and organization of the sarcomere: aberrant accumulation of one sarcomeric component at different sites in the muscle cell nucleates the subsequent recruitment of addition sarcomeric components from existing sarcomeres and/or newly made proteins. We suggest that these aggregates nucleate further addition of sarcomeric proteins, leading to nemaline bodies found in patients. Our results describing how the NM point mutations affect CFL2’s function will aid in to screening for therapeutics to improve muscle structure and function in these NM patients. Further analysis could also shed light on the contribution of muscle type, metabolism, and other pathways, to the progression of the structural changes within the muscle.

## Supporting information

Supplemental Figures 1-6 associated with main figures

## ACKNOWLEDGEMENTS

We thank the members of the Baylies lab for helpful discussions and particularly Krista Dobi for help in making the *Mhc-Gal, Zasp-GFP* stock. We also thank Silvia Jansen (Washington University) for helpful discussions and technical assistance, Kathryn Anderson, William Razzell, Jon Rosen and Krista Dobi for helpful comments on the manuscript, and our funding agencies: NIH [R21AR067361, RO1AR108981, RO1GM121971] to MKB, National Cancer Institute [P30 CA 008748] core grant to MSKCC, and NIH [R01GM063691] to BLG. We thank members of the Electron Microscopy Resource Centre at The Rockefeller University particularly Nadine Soplop and Kunihiro Uryu for technical assistance.

## AUTHOR CONTRIBUTIONS

M.B. and M.K.B. conceived and designed the study. M.B. performed all the experiments. S.F.Y. identified the phenotype. S.M.C. did all the single molecule studies and was supervised by B.L.G. D.B.S. helped with transformants and westerns. S.E.F. analyzed data. M.B. and M.K.B wrote the paper with input from all the authors. M.K.B. acquired the funding and supervised the study.

## MATERIALS AND METHODS

### Fly stocks

*Drosophila* stocks and crosses were grown on standard cornmeal medium at 25°C. For experiments using *GAL80^ts^* and *DTS5*, flies were raised at 18°C and 25°C respectively and then shifted to the permissive temperature 29°C to either inhibit GAL80^ts^ or activate DTS5 function. The *GAL4-UAS* system (Brand and Perrimon, 1993) was used for expression studies. For muscle-specific expression, *Mef2-Gal4* (Halfon et al., 2000), and *Mhc-Gal4* (Schuster et al., 1996) were used. *UAS-mCherry-RNAi* (BDSC# 35785) was used as the *control RNAi. UAS-tsr-RNAi (HMS00534)* (BDSC #65055) was used as *tsr RNAi (1)*. *UAS-tsr-IR* (VDRC #110599) was used as *tsr RNAi (2)*. *UAS-HA-PI31* (Bader et al., 2011) and *UAS-DTS5* (Bader et al., 2011) were used to manipulate the proteasome (gift of H. Stellar)*. UAS-2x-GFP* (BDSC # 6874) and *UAS-moesin::mCherry* (Millard and Martin, 2008) was used to outline the muscle to count sarcomeres. The following gene traps were used: *tsr::GFP (ZCL2393)* (DGRC #110875), *Tmod::GFP* (BDSC #50861), and *Zasp66::GFP (ZCL0663)* (BDSC #6824).

### Viability assays

Embryos were collected at 25°C on yeasted apple juice agar plates. Stage 15-16 embryos lacking the balancer (where applicable) were staged by gut morphology and hand-selected for analysis. 35 embryos were counted and transferred to a lightly yeasted apple juice agar plate and raised at 25°C overnight. The first instar larvae that hatched the following day were counted and transferred to vials of standard fly food at 25°C. The above process was repeated for 3 days. Eight to ten days later, the number of pupal cases and adults present in the vials were quantified. For each genotype the experiments were repeated twice. For the *DmCFL RNAi (1)* alone the viability experiments were repeated three times and the same values were used throughout the paper in all the graphs containing *DmCFL RNAi (1)* driven by *Mhc-Gal4*. The mean survival along with SEM from the experiments were plotted using GraphPad Prism 7.0.

### Transgenics

For the *Drosophila DmCFL^WT^* rescue experiments, *DmCFL^WT^* was PCR-amplified from EST clone LD06785 (*DGRC*\*) using the following primers: EcoRI-wt-tsr Forward: 5’ CACC GAA TTC ATG GCT TCT GGT GTA ACT GTG TCT 3’ and BglII-tsr-noSTOP Reverse : 5’ CACC GAA TTC ATG GCT TCT GGT GTA ACT ATG TCT GAT GTC TGC AAG ACT ACA T 3’. *DmCFL^V7M^* was PCR-amplified from the same EST with nucleotide 19 mutated from G to A during that amplification using the following primer EcoRI-tsr-G19A Forward: 5’ CACC GAA TTC ATG GCT TCT GGT GTA ACT ATG TCT GAT GTC TGC AAG ACT ACA T 3’ and the BglII-tsr-noSTOP Reverse primer.

The amplified products were cloned into the pENTR plasmid containing UASp and 3xHA (Life Technologies, cat# K240020), digested with EcoRI and BglII, and then ligated into pTWH (*DGRC\**, cat# 1100). All constructs were sequenced and verified constructs were injected into *w^1118^* embryos by transposable-p-element-based insertion methods (*Genetic Services*, Boston). Potential transformants were screened for *w^+^*.

For human Cofilin rescue experiments, *CFL2^wt^* was PCR-amplified from pET15b-CFL2 (Chin et al., 2016) using the following primers: EcoRI-cof-tsr-wt Forward: 5’ CACC GAA TTC ATG GCT TCT GGA GTT ACA GTG AAT and BamHI-cof2-noSTOP Reverse: 5’ GGA TCC TAA TGG TTT TCC TTC AAG TGA AAC T 3’.*CFL2^G19A^* was PCR-amplified from the same template, using a forward primer containing the desired mutation (nuc.19 G>A): 5’ CACC GAA TTC ATG GCT TCT GGA GTT ACA ATG AAT GAT GAA GTC ATC AAA GTT TTT 3’ and the BglII-Cof2-noSTOP Reverse primer.*CFL1^wt^ was* PCR-amplified from pET15b-CFL1 (Chin et al., 2016) using the following primers: EcoRI-CFL1-wt Forward: 5’ CACC GAA TTC ATG GCC TCC GGT GTG GCT GTC T 3’ and BamHI-CFL1-noSTOP Reverse: 5’ GGA TCC CAA AGG CTT GCC CTC CAG GGA G 3’.

The amplified products were cloned into the pENTR plasmid (Life Technologies, cat # K240020), digested with Eco RI and Bam HI, and then ligated into pUASg-3xHA.attB (gift from Konrad Basler, Univ. of Zurich). Constructs were injected in a site-directed fashion into attP40 (ch. 2) and attP2 (ch. 3) sites of *w^1118^* fly embryos (Rainbow Genetics). Transformants were screened for *w^+^*.

### Alignments

Protein sequence alignments were performed using MacVector 14.5.3.

### Immunostainings

Stage 15-16 embryos of the respective genotype were hand-selected and grown at 25°C for 5 days until they reached 3^rd^ instar. *DTS5-HA* flies were raised at 25°C and hand-selected, after which the first instar larvae were moved to 29°C and allowed to grow for 3 days until the larvae reached 3^rd^ instar. Larvae were dissected in ice cold HL3.1 as described (Brent et al., 2009)(Brent JR et al. 2009) and fixed with 10% formalin (Sigma, #HT501128-4L) for 20 minutes. Larval fillets were blocked with PBS supplemented with 0.1% BSA and 0.3% Triton X-100 for 30 minutes. Larval fillets were incubated with primary antibody overnight at 4°C, followed by washes in PBT-BSA and 2-hour incubation with secondary antibodies and Phalloidin at room temperature. After washes in PBT, larval fillets were then mounted in ProLong Gold (Invitrogen, #P36930). Z-stacks were acquired using either a SP5 (Leica) laser-scanning microscope with 40x/1.25NA HCX PL Apochromat oil objective or an inverted LSM-700 (Zeiss) laser-scanning confocal microscope with either a Plan-Neofluor 5x /0.16 NA, 10x/0.30NA, 20x/0.8NA air objective or a 40x/1.4NA, 63x/1.4 DiC Apochromat oil objective. Maximum intensity projections of confocal Z-stacks were rendered using Fiji (NIH). All resulting 2D projection images were cropped using Adobe Photoshop CC 2018. Approximately 8 larvae of each genotype were dissected and a minimum of 3 different images for each muscle class was acquired. The most representative image was included as a Figure. The following primary antibodies were used at the specified concentration: mouse anti-Myosin Heavy Chain (1:250, gift from S. Abmayr), chicken anti-GFP(1:200, Abcam #13970), rabbit anti-Zasp (1:400, gift from F. Schöck), rabbit anti-Lasp (1:400, gift from F.Schöck), rat anti-Tropomodulin (1:200, gift from V. Fowler), rat anti–α-actinin (1:200, Abcam #ab50599), rabbit anti-obscurin (1:200, gift from B. Bullard), rabbit anti-cpa (1:200, gift from F. Janody), rat anti-HA (1:100, from Roche #11867423001), rabbit anti-SALS (1:200, gift from N. Perrimon), mouse anti-FK2 (1:200, Enzo Lifesciences #BML-PW8810), rat anti-Troponin-T (1:200, Barbarham Institute #BT-GB1455) and rat anti-Kettin (1:200, Abcam #ab50585). Alexa Fluor 488-, 555-, and 647-conjugated fluorescent secondary antibodies (1:200), Alexa Fluor 488-, 546-, and 647-conjugated Phalloidin (1:100), and Hoechst-3342 (1μg/mL, Invitrogen) were used for fluorescent stains.

### Line profiles for protein localization

Immunofluorescent images of larval muscles were acquired using an inverted LSM-700 (Zeiss) laser-scanning confocal microscope with a 63x/1.4NA DiC Apochromat oil objective. Images were then analyzed using Fiji (NIH) where projections lines were drawn on single slice, spanning the length of 4 sarcomeres (Z disc-Z-disc) using Zasp staining. Fluorescence intensity along the lines were calculated using the Plot Profile function in Fiji (NIH). Intensity was normalized at every point along the line by using the formula: Relative fluorescence intensity= Intensity at point – Intensity minimum/ Intensity maximum X 100. A maximum of 3 lines were drawn per muscle and at least 5 different muscles from 5 different larvae were analyzed. The relative fluorescent intensities of each channel were plotted on Graph Pad Prism 7.0, to generate line profiles for protein localization.

### Larval tracking

A larva was placed at the center of 8.5 cm apple juice agar plate. 45-second movies of the larva crawling were captured on an iPhone v.6.0. Movies were converted into image sequences and uploaded onto Fiji (NIH). Each larva was manually tracked using the Manual Tracking plugin on Fiji (NIH). Average velocity was calculated by averaging velocities of the larvae over 44 frames. Approximately 8 larvae of each genotype were tracked. Genotypes were compared by student’s t-test in GraphPad Prism software.

For measuring larval velocities over different developmental stages, stage 15-16 embryos of the appropriate genotype were selected and allowed to develop at 25°C. The velocities of the larvae were determined every 24 hours, starting at the first instar stage. After tracking, the larvae were returned to the food. Approximately 8 larvae of each genotype were tracked. Velocities at different developmental stages were compared to the control larvae using ANOVA in GraphPad Prism software.

### Immunoblotting

Larval pelts for immunoblotting were generated by dissecting larva as discussed above, followed by excision of the head and tail of the larva. The remaining muscle-enriched pelts were then transferred into Larval lysis Buffer (50mM HEPES, pH 7.5, 150mM NACl, 0.5% NP40, 0.1% SDS, 2mM DTT, 1 protease inhibitor cocktail tablets (1 tablet in 10mL of lysis buffer, Roche (#11836153001)). Lysates were homogenized, and protein concentrations were determined using Bradford assay. Equal concentrations of protein (12.5μg for Tsr :GFP and 40μg for HA) were run on an either a 10% (for GFP), or a 12.5% (for HA) polyacrylamide gel after which they were transferred onto a nitrocellulose membrane (ThermoScientific, # 88018). Membranes were blocked with 5% milk in TBST (Tris-Buffered Saline +0.1% Tween) for an hour. The blots were then incubated with primary antibodies in 5% milk overnight at 4°C, and then for an hour at room temperature with secondary antibodies in 5% milk. The immunoreaction was visualized in a KwikQuant Imager (Kindle Biosciences, LLC, # D1001) using 1-Shot Digital-ECL (Kindle Biosciences, # R1003). Images were quantified using Fiji (NIH). Images were then processed using Adobe Photoshop CC 2018. Figures and quantifications are representative of 3 biological replicates and protein expression was normalized GAPDH within each sample.

The following primary antibodies were used in the specified concentrations: mouse anti-GFP (1:1000, Roche #11814460001), rat anti-HA (1:1000, Roche #11867431001) and mouse anti-GAPDH (1:10,000, Abcam #ab9484). Peroxidase-conjugated donkey anti-mouse (1:5000, Jackson Immunoresearch #715-035-151)) and Peroxidase-conjugated donkey anti-rat (1:5000, Jackson Immunoresearch #712-035-150) were used as secondary antibodies.

### Electron microscopy

Third instar larvae were dissected as described previously and were immediately placed in fixative composed of 2% glutaraldehyde, 4% Paraformaldehyde, 2mM Cacl2, 0.1% tannic acid in 0.1M sodium cacodylate buffer, pH 7.4. The samples were then fixed for 3 minutes using the Pelco Biowave (Ted Pella), followed by overnight fixation at 4°C. Samples were post-fixed with 1% osmium tetroxide containing 1.5% potassium ferrocyanide in cacodylate buffer for 1 hour on ice. They were then stained with 1% uranyl acetate aqueous solution for 30 minutes at room temperature. The samples were dehydrated in a graded series of ethanol using the Pelco Biowave and acetone dehydration at room temperature for 10 minutes. Samples were infiltrated and embedded with Eponate 12™ (Ted Pella). 60-70nm ultrathin sections were then sectioned from 3 samples for each genotype using a Reichert Jung Ultracut E microtome. Sections were stained with 2% Uranyl acetate and Sato’s lead stain and then imaged using a JEOL 1400 Plus Transmission Electron Microscope at 120kV equipped with a Gatan Ultrascan 994 US 1000XP camera and Digital Micrograph imaging software (a gift from Helmsley Charitable Trust) at the Electron Microscopy Resource Center in The Rockefeller University. For each genotype, approximately 3 biological replicates were processed and imaged. Images representative of each genotype are depicted in the Figures.

### Larval heat fixation

Larvae of the appropriate stage containing RNAi and/or GFP protein trap were analyzed as previously described (Schnorrer et al., 2010). Stage 15-16 embryos were selected and developed at 25°C. Appropriately staged larvae were fixed by brief (∼1 sec.) submersion in 65°C water and then mounted on a slide with halocarbon oil. Images were acquired using an inverted LSM-700 (Zeiss) laser-scanning confocal microscope with either a 10x/0.30NA, 20x/0.8NA air-objective. Maximum intensity projections of confocal Z-stacks were rendered using Fiii (NIH). All resulting 2D projection images were cropped using Adobe Photoshop CC 2018. For analyzing *Dm*CFL::GFP levels in *control RNAi* and *DmCFL RNAi (1)* larvae at different developmental stages, images were acquired on the same day using identical laser power and software settings.

### Time-lapse live imaging

Larvae were anesthetized as described in (Choi et al., 2014). After selecting stage 15-16 embryos of the appropriate genotype, the larvae were imaged every 24 hours from first instar onwards for 4 days. For imaging, larvae were placed on a slide with 50% glycerol and cover slip. Imaging time was limited to less than 20 minutes after which animals were washed gently with PBS, allowed to recover and returned to the food media. Images were acquired in an inverted LSM-700 (Zeiss) laser-scanning confocal microscope using the 10x/0.30NA air-objective. Maximum intensity projections of confocal Z-stacks were rendered using Fiii (NIH). All resulting 2D projection images were cropped using Adobe Photoshop CC 2018. Approximately 3 larvae of each genotype were tracked, the most representative images were included as Figures.

### Fluorescence Recovery After Photobleaching (FRAP)

*Mhc-Gal4, Zasp66::GFP* female flies were crossed to control *RNAi* or *DmCFL RNAi (1)* males. Stage 15-16 embryos were collected on apple juice-agar plates and allowed to develop for 4 days at 25°C. Third instar larvae of both genotypes were anesthetized as described (Fernandes and Schöck, 2014) by exposure for 15-20 minutes to Kwan Loong Oil (Haw Par Healthcare Ltd). Anesthetized larvae were covered in 50% glycerol and were then mounted on a glass slide with a coverslip. Heartbeat was verified before acquisition to ensure that the larvae were alive during the recording. 3 unique Regions of interest (ROIs) were selected in a single larval muscle and the Zasp66::GFP was bleached using the 488-nm laser. Fluorescence recovery of Zasp66::GFP was recorded for approximately 10 minutes with a 488-nm laser, imaging every 30 seconds. Intensity of laser was set to ensure that bleaching was not below 20-30% of initial intensity. Three muscles from three different larvae were analyzed for each genotype. Data were analyzed using Leica Application Suite X (Leica), where the intensities of the bleached, unbleached and background regions were normalized to set the prebleach intensities close to one, in order compare data across experiments. 9 independent recovery (3 biological replicates), normalized datasets were analyzed using GraphPad Prism 7.0 software. The fitted line was calculated by the software using the one-phase exponential equation. Mobile fraction for each genotype was calculated using the formula Mobile fraction (Fm)= F∞/F0. F∞ is the fluorescence intensity after full recovery and F0 is the fluorescence intensity before photobleaching. Images were acquired at room temperature with a Leica TCS SP5 II confocal microscope, using a 20x/0.75NA glycerol immersion objective.

### Muscle classes quantification

Third instar larvae of the appropriate genotypes were dissected as described above. Larval muscles stained with phalloidin as well as with a Z-disc protein were visualized using the 20x air-objective of the LSM-700 confocal microscope to classify muscles into the 3 different classes. All 30 muscles from a hemisegment were scored as either class 1,2 or 3 muscle. Only hemisegments with all 30 intact muscles were scored. A maximum of 5 hemisegments were counted per larva and 30 hemisegments in total were counted per genotype. The percentage of muscles falling into each class was calculated for each genotype. The percentages of each muscle class were calculated per genotype and plotted in GraphPad Prism 7.0, to generate graphs.

For calculating the number of muscles of different classes at each larval instar, *Tmod::GFP,Mhc-Gal4* females were crossed to control *RNAi* or *DmCFL RNAi (1)* males. Stage 15-16 embryos were collected on apple juice-agar plates and allowed to develop for at 25°C. Larvae were heat fixed every 24 hours from the first instar stage for 4 days. A maximum of 3 hemisegments per larvae, and a total of 15 hemisgements were analyzed for each developmental stage. Data were analyzed as in preceding paragraph. The percentage of each class of muscle at every larval stage was calculated. The percentages of each muscle class were plotted in GraphPad Prism 7.0, to generate graphs.

### FK2 intensity measurements

Images of VL3 and VL4 muscles from *control RNAi* and *DmCFL RNAi (1)* larvae were acquired under identical conditions using an inverted LSM-700 (Zeiss) with a 20x/0.8NA air-objective. Images were then analyzed using Fiji (NIH). First threshold intensities were set for each channel in the *control RNAi* muscles. Hoechst channel was converted to a binary image and was then subtracted from the FK2 channel, to calculate pixel intensity only in the sarcoplasm. Raw Integrated Intensity was calculated in the outlined area. Number of pixels of FK2 per µm^2^ of muscle mass was calculated by dividing the Raw Integrated Intensity with the Muscle Area. Experiment was repeated twice and plotted on GraphPad Prism 7.0 to generate a graph. P values were calculated using Student’s unpaired t-test.

### Troponin T (TnT) and Tropomodulin (Tmod) intensity measurements in the muscle

Images of VL3 and VL4 muscles from *control RNAi* and *DmCFL RNAi (1)* larvae were acquired under identical conditions using an inverted LSM-700 (Zeiss) with a 20x/0.8NA air-objective. Images were then analyzed using Fiji (NIH). Threshold intensities were initially set for TnT and Tmod channels in the *control RNAi* muscles. The center of the muscle was estimated using line tool in Fiji (NIH) and box measuring 217×68 pixels was drawn. Raw integrated pixel intensity was calculated in the outlined area. For measuring intensities for the entire muscle, the same procedure was followed as above, after threshold intensities were set the entire muscle was outlined and the raw integrated pixel intensity was calculated in the outlined area. Integrated pixel intensity of TnT/Tmod per µm^2^ of muscle mass at the center of the muscle/ entire muscle was calculated by dividing the Raw integrated pixel intensity by the area of the region measured. A minimum of 13 different Class 2 muscles from 8 different *DmCFL RNAi (1)* larvae were imaged and analyzed. Experiment was repeated twice and plotted on GraphPad Prism 7.0 to generate a graph. P values were calculated using Student’s unpaired t-test.

### Sarcomere analysis

#### Sarcomere number

Larvae of the appropriate stage containing *Zasp66::GFP (ZCL0663)* protein trap and *UAS-moesin::mCherry* driven by *Mhc-GAL4* were heatfixed as described above, and maximum intensity confocal Z-stacks were acquired. A total of 15 hemisegments from at least 5 different larvae were counted. Two different muscles VL1 and LT1 of different sizes were chosen, and number of sarcomeres in each muscle was counted by counting number of Z-lines. Numbers were plotted in GraphPad Prism 7.0, to generate graphs.

#### Muscle length and Sarcomere size

Muscle length of the above muscles were calculated by using the line function on Fiji (NIH) to draw a line from one muscle end to the other. The muscle length was then divided by the number of sarcomeres to determine average sarcomere size for each muscle at each instar. Numbers were plotted in GraphPad Prism 7.0, to generate graphs. Pearsons correlation coefficient (r) and R^2^ values were calculated using GraphPad Prism 7.0

#### Sarcomere number in Class 1 and Class 2 muscles

Early 3^rd^ instar larvae of the *Zasp66::GFP (ZCL0663)* protein trap and *UAS-moesin::mCherry* and either *UAS-mCherry RNAi/ UAS tsr TRiP RNAi* driven by *Mhc-GAL4* were heatfixed as described above, and maximum intensity confocal Z-stacks were acquired. A total of 9 hemisegments from 5 different larvae containing one Class 1 VL muscle and a Class 2 were counted by counting number of moesin-mCherry aggregates. Pearsons correlation coefficient (r) and R^2^ values were calculated using GraphPad Prism 7.0. Number of sarcomeres were plotted against muscle length in GraphPad Prism 7.0, to generate graphs.

#### Plasmids and protein purification

The plasmid for *E. coli* expression of human Cof2 was a gift from Dr. David Kovar (University of Chicago). Site-directed mutagenesis was performed on this plasmid to introduce the V7M mutation, and the resulting plasmid was verified by DNA sequencing. Rabbit skeletal muscle actin was purified, and in some preparations labeled on Cys^374^ with Oregon Green maleimide (Life Technologies), as described in detail (Graziano et al., 2013). Human CFL2 and Cof2^V7M^ were purified from *E. coli* as described (Chin et al., 2016). In brief, CFL2 and CFL2^V7M^ were expressed in BL21 (DE3) *E. Coli*, grown to log phase at 37⁰C in TB medium, and expression was induced with 1 mM isopropyl beta-D-1-thiogalactopyranoside at 18⁰C for 15 hr. Cells were harvested by centrifugation and stored as pellets at −80⁰C. Cell pellets were resuspended in 20 mM Tris pH 8.0, 50 mM NaCl, 1 mM DTT, and protease inhibitors, then lysed by sonication. The lysate was cleared at 30,000 *g* for 30 min in a Fiberlite F13-14X50CY rotor (Thermo Fisher Scientific), then loaded on a 5 ml HiTrap HP Q column (GE Healthcare Biosciences). The flow-through, containing Cof2^WT^ or Cof2^V7M^, was dialyzed against 20 mM Hepes pH 6.8, 25 mM NaCl, and 1 mM DTT, loaded on a 5 ml HiTrap SP FF column (GE Healthcare Biosciences), and eluted with a linear gradient of NaCl (20– 500 mM). Peak fractions were concentrated and dialyzed against 20 mM Tris pH 8.0, 50 mM KCl, and 1 mM DTT, aliquoted, and stored at −80°C.

#### Nucleotide exchange assays

To prepare ADP-G-actin, monomeric ATP-bound 2 μM rabbit muscle actin was treated overnight at 4°C with analytical grade anion exchange resin (BioRad), hexokinase (Sigma-Aldrich), and excess ADP. To initiate a reaction, 2 μM ADP-bound actin monomers and the indicated concentration of Cof2 or Cof2 ^V7M^ in CDT buffer (0.2 mM CaCl2, 0.2 mM DTT, 10 mM Tris pH 8.0), or buffer alone, were mixed and added to 50 μM ε-ATP. A fluorescence spectrophotometer (Photon Technology International) was used to monitor the reaction at 350-nm excitation and 410-nm emission at 25°C for 200 s.

#### TIRF microscopy

To prepare slides, 24×60 mm coverslips (Thermo Fisher Scientific) were cleaned by successive sonications as follows: 60 min in detergent, 20 min in 1 M KOH, 20 min in 1 M HCl min, and 60 min in ethanol. Next, coverslips were thoroughly washed with ddH2O, then dried using an N2-stream. Then, each slide was layered with 200 μl of a solution consisting of 80% ethanol pH 2.0, 2 mg/ml methoxy-poly (ethylene glycol)-silane and 2 μg/ml biotin-poly (ethylene glycol)-silane (Laysan Bio Inc.), and incubated for 16 h at 70°C. To assemble flow cells, PEG-coated coverslips were thoroughly rinsed with ddH2O and dried using an N2-stream, then attached to a prepared flow chamber (Ibidi) with double sided tape (2.5 cm × 2 mm × 120 μm) and five min epoxy resin. Immediately prior to use, coverslips were subjected to sequential incubations as follows: 3 min in HEK-BSA (20 mM Hepes pH 7.5, 1 mM EDTA, 50 mM KCl, 1% BSA), 30 s in Streptavidin (0.1 mg/ml in PBS), a fast rinse in HEK-BSA, and then equilibration in 1X TIRF buffer (10 mM imidazole, 50 mM KCl, 1 mM MgCl2, 1 mM EGTA, 0.2 mM ATP, 10 mM DTT, 15 mM glucose, 20 μg/ml catalase, 100 μg/ml glucose oxidase, and 0.5% methylcellulose (4000 cP), pH 7.5).

To measure effects on filament severing, actin monomers (10% OG-labeled, 0.5% biotinylated) were first diluted to 1 μM in TIRF buffer, and immediately transferred to a flow chamber. Actin was polymerized at room temperature until filaments reached approximately 10–15 μm in length, then free actin monomers were washed out, and TIRF buffer containing CFL2 or CFL2 ^V7M^ were introduced by flow in. Time-lapse TIRF microscopy was performed using a Nikon-Ti200 inverted microscope equipped with a 150 mW Ar-Laser (Mellot Griot), a TIRF-objective with a N.A. of 1.49 (Nikon Instruments Inc.), and an EMCCD camera (Andor Ixon). Optimal focus was maintained throughout the recordings using the perfect focus system (Nikon Instruments Inc.). Pixel size, 0.27 μm. TIRF data was analyzed using ImageJ software. The background was subtracted prior to analysis using the background subtraction tool (rolling ball radius = 50 pixels).

## REFERENCES

Abe, H., Obinata, T., 1989. An actin-depolymerizing protein in embryonic chicken skeletal muscle: purification and characterization. J. Biochem. 106, 172–180. doi:10.1093/oxfordjournals.jbchem.a122810

Abe, H., Ohshima, S., Obinata, T., 1989. A cofilin-like protein is involved in the regulation of actin assembly in developing skeletal muscle. J. Biochem. 106, 696–702. doi:10.1093/oxfordjournals.jbchem.a122919

Agrawal, P.B., Greenleaf, R.S., Tomczak, K.K., Lehtokari, V.-L., Wallgren-Pettersson, C., Wallefeld, W., Laing, N.G., Darras, B.T., Maciver, S.K., Dormitzer, P.R., Beggs, A.H., 2007. Nemaline myopathy with minicores caused by mutation of the CFL2 gene encoding the skeletal muscle actin-binding protein, cofilin-2. Am. J. Hum. Genet. 80, 162–167. doi:10.1086/510402

Agrawal, P.B., Joshi, M., Savic, T., Chen, Z., Beggs, A.H., 2012. Normal myofibrillar development followed by progressive sarcomeric disruption with actin accumulations in a mouse Cfl2 knockout demonstrates requirement of cofilin-2 for muscle maintenance. Hum. Mol. Genet. 21, 2341–2356. doi:10.1093/hmg/dds053

Agrawal, P.B., Strickland, C.D., Midgett, C., Morales, A., Newburger, D.E., Poulos, M.A., Tomczak, K.K., Ryan, M.M., Iannaccone, S.T., Crawford, T.O., Laing, N.G., Beggs, A.H., 2004. Heterogeneity of nemaline myopathy cases with skeletal muscle alpha-actin gene mutations. Ann. Neurol. 56, 86–96. doi:10.1002/ana.20157

Andrianantoandro, E., Pollard, T.D., 2006. Mechanism of actin filament turnover by severing and nucleation at different concentrations of ADF/cofilin. Mol. Cell 24, 13–23. doi:10.1016/j.molcel.2006.08.006

Bachman, J.F., Klose, A., Liu, W., Paris, N.D., Blanc, R.S., Schmalz, M., Knapp, E., Chakkalakal, J.V., 2018. Prepubertal skeletal muscle growth requires Pax7-expressing satellite cell-derived myonuclear contribution. Development 145, dev167197. doi:10.1242/dev.167197

Bader, M., Benjamin, S., Wapinski, O.L., Smith, D.M., Goldberg, A.L., Steller, H., 2011. A conserved F box regulatory complex controls proteasome activity in Drosophila. Cell 145, 371–382. doi:10.1016/j.cell.2011.03.021

Bai, J., Hartwig, J.H., Perrimon, N., 2007. SALS, a WH2-domain-containing protein, promotes sarcomeric actin filament elongation from pointed ends during Drosophila muscle growth. Dev. Cell 13, 828–842. doi:10.1016/j.devcel.2007.10.003

Bamburg, J.R., Bray, D., 1987. Distribution and cellular localization of actin depolymerizing factor. J. Cell Biol. 105, 2817–2825.

Bate, M., 1990. The embryonic development of larval muscles in Drosophila. Development 110, 791–804.

Belote, J.M., Fortier, E., 2002. Targeted expression of dominant negative proteasome mutants in Drosophila melanogaster. Genesis 34, 80–82. doi:10.1002/gene.10131

Blair, A., Tomlinson, A., Pham, H., Gunsalus, K.C., Goldberg, M.L., Laski, F.A., 2006. Twinstar, the Drosophila homolog of cofilin/ADF, is required for planar cell polarity patterning. Development 133, 1789–1797. doi:10.1242/dev.02320

Brand, A.H., Perrimon, N., 1993. Targeted gene expression as a means of altering cell fates and generating dominant phenotypes. Development 118, 401–415.

Brent, J.R., Werner, K.M., McCabe, B.D., 2009. Drosophila larval NMJ dissection. JoVE. doi:10.3791/1107

Cassandrini, D., Trovato, R., Rubegni, A., Lenzi, S., Fiorillo, C., Baldacci, J., Minetti, C., Astrea, G., Bruno, C., Santorelli, F.M., Italian Network on Congenital Myopathies, 2017. Congenital myopathies: clinical phenotypes and new diagnostic tools. Ital J Pediatr 43, 101. doi:10.1186/s13052-017-0419-z

Chin, S.M., Jansen, S., Goode, B.L., 2016. TIRF microscopy analysis of human Cof1, Cof2, and ADF effects on actin filament severing and turnover. J. Mol. Biol. 428, 1604–1616. doi:10.1016/j.jmb.2016.03.006

Choi, B.J., Imlach, W.L., Jiao, W., Wolfram, V., Wu, Y., Grbic, M., Cela, C., Baines, R.A., Nitabach, M.N., McCabe, B.D., 2014. Miniature neurotransmission regulates Drosophila synaptic structural maturation. Neuron 82, 618–634. doi:10.1016/j.neuron.2014.03.012

Demontis, F., Perrimon, N., 2009. Integration of Insulin receptor/Foxo signaling and dMyc activity during muscle growth regulates body size in Drosophila. Development 136, 983–993. doi:10.1242/dev.027466

Dix, D.J., Eisenberg, B.R., 1990a. Myosin mRNA accumulation and myofibrillogenesis at the myotendinous junction of stretched muscle fibers. J. Cell Biol. 111, 1885–1894. doi:10.1083/jcb.111.5.1885

Dix, D.J., Eisenberg, B.R., 1990b. Myosin mRNA accumulation and myofibrillogenesis at the myotendinous junction of stretched muscle fibers. J. Cell Biol. 111, 1885–1894. doi:10.1083/jcb.111.5.1885

Edwards, K.A., Montague, R.A., Shepard, S., Edgar, B.A., Erikson, R.L., Kiehart, D.P., 1994. Identification of Drosophila cytoskeletal proteins by induction of abnormal cell shape in fission yeast. Proc. Natl. Acad. Sci. U.S.A. 91, 4589–4593.

Farah, C.S., Reinach, F.C., 1995. The troponin complex and regulation of muscle contraction. FASEB J. 9, 755–767.

Fernandes, I., Schöck, F., 2014. The nebulin repeat protein Lasp regulates I-band architecture and filament spacing in myofibrils. J. Cell Biol. 206, 559–572. doi:10.1083/jcb.201401094

Fowler, V.M., 1996. Regulation of actin filament length in erythrocytes and striated muscle. Curr. Opin. Cell Biol. 8, 86–96.

Graziano, B.R., Jonasson, E.M., Pullen, J.G., Gould, C.J., Goode, B.L., 2013. Ligand-induced activation of a formin-NPF pair leads to collaborative actin nucleation. J. Cell Biol. 201, 595–611. doi:10.1083/jcb.201212059

Gunsalus, K.C., Bonaccorsi, S., Williams, E., Verni, F., Gatti, M., Goldberg, M.L., 1995. Mutations in twinstar, a Drosophila gene encoding a cofilin/ADF homologue, result in defects in centrosome migration and cytokinesis. J. Cell Biol. 131, 1243–1259.

Gupta, V.A., Beggs, A.H., 2014. Kelch proteins: emerging roles in skeletal muscle development and diseases. Skelet Muscle 4, 11. doi:10.1186/2044-5040-4-11

Gurniak, C.B., Chevessier, F., Jokwitz, M., Jönsson, F., Perlas, E., Richter, H., Matern, G., Boyl, P.P., Chaponnier, C., Fürst, D., Schröder, R., Witke, W., 2014a. Severe protein aggregate myopathy in a knockout mouse model points to an essential role of cofilin2 in sarcomeric actin exchange and muscle maintenance. Eur. J. Cell Biol. 93, 252–266. doi:10.1016/j.ejcb.2014.01.007

Gurniak, C.B., Chevessier, F., Jokwitz, M., Jönsson, F., Perlas, E., Richter, H., Matern, G., Boyl, P.P., Chaponnier, C., Fürst, D., Schröder, R., Witke, W., 2014b. Severe protein aggregate myopathy in a knockout mouse model points to an essential role of cofilin2 in sarcomeric actin exchange and muscle maintenance. Eur. J. Cell Biol. 93, 252–266. doi:10.1016/j.ejcb.2014.01.007

Haas, J.N., 1950. Cytoplasmic growth in the muscle fibers of larvae of Drosophila melanogaster. Growth 14, 277–294.

Halfon, M.S., Carmena, A., Gisselbrecht, S., Sackerson, C.M., Jiménez, F., Baylies, M.K., Michelson, A.M., 2000. Ras pathway specificity is determined by the integration of multiple signal-activated and tissue-restricted transcription factors. Cell 103, 63–74.

Henderson, C.A., Gomez, C.G., Novak, S.M., Mi-Mi, L., Gregorio, C.C., 2017. Overview of the Muscle Cytoskeleton. Compr Physiol 7, 891–944. doi:10.1002/cphy.c160033

Ilkovski, B., Cooper, S.T., Nowak, K., Ryan, M.M., Yang, N., Schnell, C., Durling, H.J., Roddick, L.G., Wilkinson, I., Kornberg, A.J., Collins, K.J., Wallace, G., Gunning, P., Hardeman, E.C., Laing, N.G., North, K.N., 2001. Nemaline myopathy caused by mutations in the muscle alpha-skeletal-actin gene. Am. J. Hum. Genet. 68, 1333–1343. doi:10.1086/320605

Jockusch, B.M., Veldman, H., Griffiths, G.W., van Oost, B.A., Jennekens, F.G., 1980. Immunofluorescence microscopy of a myopathy. alpha-Actinin is a major constituent of nemaline rods. Exp. Cell Res. 127, 409–420.

Kaya-Çopur, A., Schnorrer, F., 2019. RNA Interference Screening for Genes Regulating Drosophila Muscle Morphogenesis. Methods Mol. Biol. 1889, 331–348. doi:10.1007/978-1-4939-8897-6_20

Kondo, E., Nishimura, T., Kosho, T., Inaba, Y., Mitsuhashi, S., Ishida, T., Baba, A., Koike, K., Nishino, I., Nonaka, I., Furukawa, T., Saito, K., 2012. Recessive RYR1 mutations in a patient with severe congenital nemaline myopathy with ophthalomoplegia identified through massively parallel sequencing. Am. J. Med. Genet. A 158A, 772–778. doi:10.1002/ajmg.a.35243

Kremneva, E., Makkonen, M.H., Skwarek-Maruszewska, A., Gateva, G., Michelot, A., Dominguez, R., Lappalainen, P., 2014a. Cofilin-2 controls actin filament length in muscle sarcomeres. Dev. Cell 31, 215–226. doi:10.1016/j.devcel.2014.09.002

Kremneva, E., Makkonen, M.H., Skwarek-Maruszewska, A., Gateva, G., Michelot, A., Dominguez, R., Lappalainen, P., 2014b. Cofilin-2 controls actin filament length in muscle sarcomeres. Dev. Cell 31, 215–226. doi:10.1016/j.devcel.2014.09.002

Lexell, J., Sjöström, M., Nordlund, A.S., Taylor, C.C., 1992. Growth and development of human muscle: a quantitative morphological study of whole vastus lateralis from childhood to adult age. Muscle Nerve 15, 404–409. doi:10.1002/mus.880150323

Littlefield, R.S., Fowler, V.M., 2008. Thin filament length regulation in striated muscle sarcomeres: pointed-end dynamics go beyond a nebulin ruler. Semin. Cell Dev. Biol. 19, 511–519. doi:10.1016/j.semcdb.2008.08.009

Maggi, L., Scoto, M., Cirak, S., Robb, S.A., Klein, A., Lillis, S., Cullup, T., Feng, L., Manzur, A.Y., Sewry, C.A., Abbs, S., Jungbluth, H., Muntoni, F., 2013. Congenital myopathies--clinical features and frequency of individual subtypes diagnosed over a 5-year period in the United Kingdom. Neuromuscul. Disord. 23, 195–205. doi:10.1016/j.nmd.2013.01.004

Malfatti, E., Lehtokari, V.-L., Böhm, J., de Winter, J.M., Schäffer, U., Estournet, B., Quijano-Roy, S., Monges, S., Lubieniecki, F., Bellance, R., Viou, M.T., Madelaine, A., Wu, B., Taratuto, A.L., Eymard, B., Pelin, K., Fardeau, M., Ottenheijm, C.A.C., Wallgren-Pettersson, C., Laporte, J., Romero, N.B., 2014. Muscle histopathology in nebulin-related nemaline myopathy: ultrastrastructural findings correlated to disease severity and genotype. Acta Neuropathol Commun 2, 44. doi:10.1186/2051-5960-2-44

Malfatti, E., Romero, N.B., 2016. Nemaline myopathies: State of the art. Rev. Neurol. (Paris) 172, 614–619. doi:10.1016/j.neurol.2016.08.004

Mardahl-Dumesnil, M., Fowler, V.M., 2001. Thin filaments elongate from their pointed ends during myofibril assembly in Drosophila indirect flight muscle. J. Cell Biol. 155, 1043–1053. doi:10.1083/jcb.200108026

Michelot, A., Berro, J., Guérin, C., Boujemaa-Paterski, R., Staiger, C.J., Martiel, J.-L., Blanchoin, L., 2007. Actin-filament stochastic dynamics mediated by ADF/cofilin. Curr. Biol. 17, 825–833. doi:10.1016/j.cub.2007.04.037

Millard, T.H., Martin, P., 2008. Dynamic analysis of filopodial interactions during the zippering phase of Drosophila dorsal closure. Development 135, 621–626. doi:10.1242/dev.014001

Miyatake, S., Mitsuhashi, S., Hayashi, Y.K., Purevjav, E., Nishikawa, A., Koshimizu, E., Suzuki, M., Yatabe, K., Tanaka, Y., Ogata, K., Kuru, S., Shiina, M., Tsurusaki, Y., Nakashima, M., Mizuguchi, T., Miyake, N., Saitsu, H., Ogata, K., Kawai, M., Towbin, J., Nonaka, I., Nishino, I., Matsumoto, N., 2017. Biallelic Mutations in MYPN, Encoding Myopalladin, Are Associated with Childhood-Onset, Slowly Progressive Nemaline Myopathy. Am. J. Hum. Genet. 100, 169–178. doi:10.1016/j.ajhg.2016.11.017

Ng, J., Luo, L., 2004. Rho GTPases regulate axon growth through convergent and divergent signaling pathways. Neuron 44, 779–793. doi:10.1016/j.neuron.2004.11.014

Nilipour, Y., Nafissi, S., Tjust, A.E., Ravenscroft, G., Hossein-Nejad Nedai, H., Taylor, R., Varasteh, V., Pedrosa Domellöf, F., Zangi, M., Tonekaboni, S.H., Olivé, M., Kiiski, K., Sagath, L., Davis Laing, N., Tajsharghi, H., 2018. Ryanodine receptor type 3 (RYR3) as a novel gene associated with a myopathy with nemaline bodies. Eur. J. Neurol. doi:10.1111/ene.13607

Nworu, C.U., Kraft, R., Schnurr, D.C., Gregorio, C.C., Krieg, P.A., 2015. Leiomodin 3 and tropomodulin 4 have overlapping functions during skeletal myofibrillogenesis. J. Cell. Sci. 128, 239–250. doi:10.1242/jcs.152702

Ockeloen, C.W., Gilhuis, H.J., Pfundt, R., Kamsteeg, E.J., Agrawal, P.B., Beggs, A.H., Dara Hama-Amin, A., Diekstra, A., Knoers, N.V.A.M., Lammens, M., van Alfen, N., 2012. Congenital myopathy caused by a novel missense mutation in the CFL2 gene. Neuromuscul. Disord. 22, 632–639. doi:10.1016/j.nmd.2012.03.008

Ong, R.W., AlSaman, A., Selcen, D., Arabshahi, A., Yau, K.S., Ravenscroft, G., Duff, R.M., Atkinson, V., Allcock, R.J., Laing, N.G., 2014. Novel cofilin-2 (CFL2) four base pair deletion causing nemaline myopathy. J. Neurol. Neurosurg. Psychiatr. 85, 1058–1060. doi:10.1136/jnnp-2014-307608

Ono, K., Parast, M., Alberico, C., Benian, G.M., Ono, S., 2003. Specific requirement for two ADF/cofilin isoforms in distinct actin-dependent processes in Caenorhabditis elegans. J. Cell. Sci. 116, 2073–2085. doi:10.1242/jcs.00421

Ono, S., 2010. Dynamic regulation of sarcomeric actin filaments in striated muscle. Cytoskeleton (Hoboken) 67, 677–692. doi:10.1002/cm.20476

Ono, S., Baillie, D.L., Benian, G.M., 1999. UNC-60B, an ADF/cofilin family protein, is required for proper assembly of actin into myofibrils in Caenorhabditis elegans body wall muscle. J. Cell Biol. 145, 491–502.

Pearson, A.M., 1990. Muscle growth and exercise. Crit Rev Food Sci Nutr 29, 167–196. doi:10.1080/10408399009527522

Romero, N.B., Sandaradura, S.A., Clarke, N.F., 2013. Recent advances in nemaline myopathy. Curr. Opin. Neurol. 26, 519–526. doi:10.1097/WCO.0b013e328364d681

Sandaradura, S.A., Bournazos, A., Mallawaarachchi, A., Cummings, B.B., Waddell, L.B., Jones, K.J., Troedson, C., Sudarsanam, A., Nash, B.M., Peters, G.B., Algar, E.M., MacArthur, D.G., North, K.N., Brammah, S., Charlton, A., Laing, N.G., Wilson, M.J., Davis, M.R., Cooper, S.T., 2018. Nemaline myopathy and distal arthrogryposis associated with an autosomal recessive TNNT3 splice variant. Human Mutation 39, 383–388. doi:10.1002/humu.23385

Sanger, J.M., Sanger, J.W., 2008. The dynamic Z bands of striated muscle cells. Sci Signal 1, pe37–pe37. doi:10.1126/scisignal.132pe37

Sanoudou, D., Beggs, A.H., 2001. Clinical and genetic heterogeneity in nemaline myopathy – a disease of skeletal muscle thin filaments. Trends in Molecular Medicine 7, 362–368. doi:10.1016/S1471-4914(01)02089-5

Schnorrer, F., Schönbauer, C., Langer, C.C.H., Dietzl, G., Novatchkova, M., Schernhuber, K., Fellner, M., Azaryan, A., Radolf, M., Stark, A., Keleman, K., Dickson, B.J., 2010. Systematic genetic analysis of muscle morphogenesis and function in Drosophila. Nature 464, 287–291. doi:10.1038/nature08799

Schuster, C.M., Davis, G.W., Fetter, R.D., Goodman, C.S., 1996. Genetic dissection of structural and functional components of synaptic plasticity. I. Fasciclin II controls synaptic stabilization and growth. Neuron 17, 641–654.

Sevdali, M., Kumar, V., Peckham, M., Sparrow, J., 2013. Human congenital myopathy actin mutants cause myopathy and alter Z-disc structure in Drosophila flight muscle. Neuromuscul. Disord. 23, 243–255. doi:10.1016/j.nmd.2012.11.013

Sewry, C.A., Laitila, J.M., Wallgren-Pettersson, C., 2019. Nemaline myopathies: a current view. J. Muscle Res. Cell. Motil. 53, 564–16. doi:10.1007/s10974-019-09519-9

Singh, S.H., Kumar, P., Ramachandra, N.B., Nongthomba, U., 2014. Roles of the troponin isoforms during indirect flight muscle development in Drosophila. J. Genet. 93, 379–388.

Stevenson, T.O., Mercer, K.B., Cox, E.A., Szewczyk, N.J., Conley, C.A., Hardin, J.D., Benian, G.M., 2007. unc-94 encodes a tropomodulin in Caenorhabditis elegans. J. Mol. Biol. 374, 936–950. doi:10.1016/j.jmb.2007.10.005

Subramanian, K., Gianni, D., Balla, C., Assenza, G.E., Joshi, M., Semigran, M.J., Macgillivray, T.E., Van Eyk, J.E., Agnetti, G., Paolocci, N., Bamburg, J.R., Agrawal, P.B., Del Monte, F., 2015. Cofilin-2 phosphorylation and sequestration in myocardial aggregates: novel pathogenetic mechanisms for idiopathic dilated cardiomyopathy. J. Am. Coll. Cardiol. 65, 1199–1214. doi:10.1016/j.jacc.2015.01.031

Sztal, T.E., Zhao, M., Williams, C., Oorschot, V., Parslow, A.C., Giousoh, A., Yuen, M., Hall, T.E., Costin, A., Ramm, G., Bird, P.I., Busch-Nentwich, E.M., Stemple, D.L., Currie, P.D., Cooper, S.T., Laing, N.G., Nowak, K.J., Bryson-Richardson, R.J., 2015. Zebrafish models for nemaline myopathy reveal a spectrum of nemaline bodies contributing to reduced muscle function. Acta Neuropathol. 130, 389–406. doi:10.1007/s00401-015-1430-3.

Thirion, C., Stucka, R., Mendel, B., Gruhler, A., Jaksch, M., Nowak, K. J., Binz, N., Laing, N. G. and Lochmüller, H. (2001). Characterization of human muscle type cofilin (CFL2) in normal and regenerating muscle. Eur. J. Biochem. 268, 3473–3482.

Vartiainen, M.K., Mustonen, T., Mattila, P.K., Ojala, P.J., Thesleff, I., Partanen, J., Lappalainen, P., 2002. The three mouse actin-depolymerizing factor/cofilins evolved to fulfill cell-type-specific requirements for actin dynamics. Mol. Biol. Cell 13, 183–194. doi:10.1091/mbc.01-07-0331

Viswanathan, M.C., Blice-Baum, A.C., Schmidt, W., Foster, D.B., Cammarato, A., 2015. Pseudo acetylation of K326 and K328 of actin disrupts Drosophila melanogaster indirect flight muscle structure and performance. Front Physiol 6, 116. doi:10.3389/fphys.2015.00116

Wallgren-Pettersson, C., Jasani, B., Newman, G.R., Morris, G.E., Jones, S., Singhrao, S., Clarke, A., Virtanen, I., Holmberg, C., Rapola, J., 1995. Alpha-actinin in nemaline bodies in congenital nemaline myopathy: immunological confirmation by light and electron microscopy. Neuromuscul. Disord. 5, 93–104.

Wallgren-Pettersson, C., Pelin, K., Hilpelä, P., Donner, K., Porfirio, B., Graziano, C., Swoboda, K.J., Fardeau, M., Urtizberea, J.A., Muntoni, F., Sewry, C., Dubowitz, V., Iannaccone, S., Minetti, C., Pedemonte, M., Seri, M., Cusano, R., Lammens, M., Castagna-Sloane, A., Beggs, A.H., Laing, N.G., la Chapelle, de A., 1999. Clinical and genetic heterogeneity in autosomal recessive nemaline myopathy. Neuromuscul. Disord. 9, 564–572.

Wallgren-Pettersson, C., Pelin, K., Nowak, K.J., Muntoni, F., Romero, N.B., Goebel, H.H., North, K.N., Beggs, A.H., Laing, N.G., ENMC International Consortium On Nemaline Myopathy, 2004. Genotype-phenotype correlations in nemaline myopathy caused by mutations in the genes for nebulin and skeletal muscle alpha-actin. Neuromuscul. Disord. 14, 461–470. doi:10.1016/j.nmd.2004.03.006

Yamaguchi, M., Robson, R.M., Stromer, M.H., Dahl, D.S., Oda, T., 1982. Nemaline myopathy rod bodies. Structure and composition. J. Neurol. Sci. 56, 35–56.

Yang, H., Schmidt, L.P., Wang, Z., Yang, X., Shao, Y., Borg, T.K., Markwald, R., Runyan, R., Gao, B.Z., 2016. Dynamic Myofibrillar Remodeling in Live Cardiomyocytes under Static Stretch. Sci Rep 6, 20674. doi:10.1038/srep20674

Zhang, S., Bernstein, S.I., 2001. Spatially and temporally regulated expression of myosin heavy chain alternative exons during Drosophila embryogenesis. Mech. Dev. 101, 35–45.

