## Supplemental Figures 1-6 associated with main figures for "Cofilin loss in *Drosophila* contributes to myopathy through defective sarcomerogenesis and aggregate formation during muscle growth"

*Balakrishnan et al., Supplementary Figure 1 - Supplement to Figure 1*

### Supplementary Figure 1: Muscle-specific knockdown of *Dm*CFL affects muscle structure and function

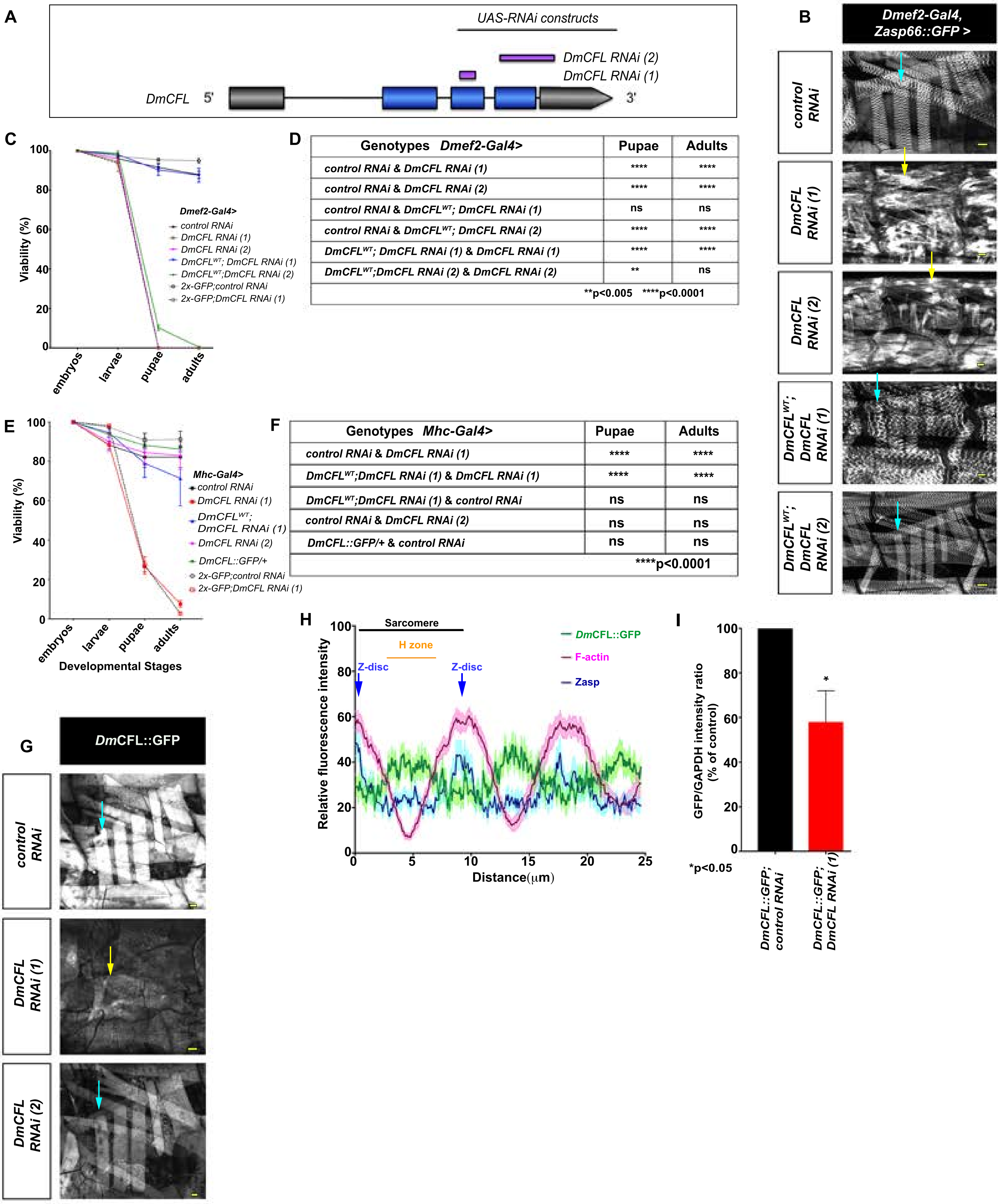


D

| Genotypes Dmef2-Gal4> | Pupae | Adults |
|---|---|---|
| control RNAi & DmCFL RNAi (1) | **** | **** |
| control RNAi & DmCFL RNAi (2) | **** | **** |
| control RNAi & DmCFL$^{WT}$; DmCFL RNAi (1) | ns | ns |
| control RNAi & DmCFL$^{WT}$; DmCFL RNAi (2) | **** | **** |
| DmCFL$^{WT}$; DmCFL RNAi (1) & DmCFL RNAi (1) | **** | **** |
| DmCFL$^{WT}$;DmCFL RNAi (2) & DmCFL RNAi (2) | ** | ns |
| **p<0.005 ****p<0.0001 | | |

F

| Genotypes Mhc-Gal4> | Pupae | Adults |
|---|---|---|
| control RNAi & DmCFL RNAi (1) | **** | **** |
| DmCFL$^{WT}$;DmCFL RNAi (1) & DmCFL RNAi (1) | **** | **** |
| DmCFL$^{WT}$;DmCFL RNAi (1) & control RNAi | ns | ns |
| control RNAi & DmCFL RNAi (2) | ns | ns |
| DmCFL::GFP/+ & control RNAi | ns | ns |
| ****p<0.0001 | | |

***Balakrishnan et al., Supplementary Figure 1 Legend***

**Supplementary Figure 1: Muscle-specific knockdown of *Dm*CFL affects muscle structure and function – Supplement to Figure 1**

A. Schematic of *DmCFL* locus in *Drosophila*. Purple bars indicate region targeted by *UAS-DmCFL RNAi* constructs. Grey bars, untranslated regions. Blue bars, exons. Black lines, introns.

B. Larval muscle cells at 2nd instar stage expressing the listed UAS-RNAi constructs driven by *DMef2-Gal4*, an early embryonic muscle Gal4 driver. Several individual muscles from one representative hemisegment are shown. Sarcomeres are defined by expression of the Z-disc protein, Zasp-GFP (greyscale). Control muscles (top panel, one example muscle – blue arrow) have a repetitive sarcomere organization (repeated grey lines). In *DmCFL RNAi (1)* and *DmCFL RNAi (2)* muscles (second and third panels, respectively), sarcomeres are disrupted, and large ZASP-positive aggregates are formed (examples, yellow arrows). Expression of $DmCFL^{WT}$ with both the *DmCFL RNAi*'s (fourth and fifth panels) rescues the muscle phenotypes (eg. blue arrows). Images are from one of two independent experiments with N=5 larvae of each genotype at each instar.

C. Viability of flies expressing *control RNAi*, *DmCFL RNAi (1), DmCFL RNAi (2),* $DmCFL^{wt}$*; DmCFL RNAi (1),* $DmCFL^{wt}$*; DmCFL RNAi (2), 2x-GFP; control RNAi, or 2x-GFP, DmCFL RNAi (1)* constructs using *Dmef2-Gal4.* All of the *DmCFL RNAi (1)* and *DmCFL RNAi (2)* larvae die at the 2nd larval instar stage. Expression of $DmCFL^{WT}$ in combination with *DmCFL RNAi (1*) rescues the viability of the organism significantly, increasing both the number of pupae as well as adults that eclose. Expression of $DmCFL^{WT}$ in combination with *DmCFL RNAi (2*) partially rescues the viability of the organism, increasing the number of pupae. The rescue of both *DmCFL RNAi*'s with $DmCFL^{WT}$ suggests that *DmCFL* expression in the muscle is necessary for survival of the organism. N=100 embryos of each genotype were tracked over the different developmental stages. No significant difference in the viability of *control RNAi*, vs. *2x-GFP; control RNAi* and *DmCFL RNAi (1) vs. 2x-GFP,DmCFL RNAi (1)* constructs. Error bars, mean±SEM.

D. Statistical analysis performed on the viability of the different stages of development (panel C). There was no significant difference in the viability of the genotypes at the embryo and larval stages, only at later stages of development. P-values calculated by 2-way ANOVA test.

E. Viability of flies expressing *control RNAi*, *DmCFL RNAi (1), DmCFL RNAi (2),*or *DmCFL$^{wt}$; DmCFL RNAi (1), 2x-GFP; control RNAi, or 2x-GFP,DmCFL RNAi (1)* constructs using *Mhc-Gal4,* a later embryonic muscle Gal4 driver. The majority of *DmCFL RNAi (1)* larvae fail to pupate and die at the end of 3$^{rd}$ instar. Expression of *DmCFL$^{WT}$* with *DmCFL RNAi (1*) partially rescues the viability of the organism, increasing both the number of pupae as well as adults that eclose, suggesting that *DmCFL* expression in the muscle cell is necessary for survival of the organism. Expression of *DmCFL RNAi (2)* driven by *Mhc-Gal4* did not affect viability of the organism. No significant difference in the viability of *control RNAi*, vs. *2x-GFP; control RNAi* and *DmCFL RNAi (1) vs. 2x-GFP,DmCFL RNAi (1)* constructs.

F. Statistical analysis performed on the viability of the different stages of development (panel E). There was no significant difference in the viability of the genotypes at the embryo and larval stages. P-values calculated by 2-way ANOVA test.

G. Larval muscles at 3$^{rd}$ instar stage expressing *Dm*CFL::GFP (greyscale) with *control RNAi*, *DmCFL RNAi (1),* or *DmCFL RNAi (2*) using *Mhc-Gal4* as a driver. In the presence of *DmCFL RNAi (1), Dm*CFL::GFP levels are drastically reduced (eg. yellow arrow). However, in the presence of *DmCFL RNAi (2), Dm*CFL::GFP is not completely lost (eg. blue arrow). This expression level could explain the lack of muscle phenotypes found in these animals. Representative images are from one of three independent experiments with N=5 larvae of each genotype. All images were taken with the same microscope settings.

H. Relative average fluorescence intensities in the absence of a GFP secondary antibody of endogenous *Dm*CFL-GFP (green), F-actin (magenta), and Zasp (blue) across the length of 4 sarcomeres (average ± SEM). *Dm*CFL::GFP localizes to the Z-disc of the sarcomere as well as the F-actin free region of the sarcomere, i.e. the H-zone, as seen in using GFP secondary antibody in Figure 1G. N = 10 measurements from at least 5 different muscles from 5 larvae.

I. *Dm*CFL-GFP protein levels in *control RNAi* or *DmCFL RNAi (1)* dissected larval carcasses, which include muscles, epidermis, and sensory neurons. GFP levels were normalized to GAPDH levels in each sample in 3 independent experiments with N=20 larvae per genotype. * $p < 0.05$. P values by Unpaired *t*-test. Error bars, mean±SEM. We observed a 60% knockdown relative to control levels in total animal carcasses. A more significant reduction in *Dm*CFL levels would be observed if we could isolate body muscles only, as suggested by IF in S1G. The isolation of larval muscles is not technically feasible at this time.

Scale Bar 25μm (B, G)

### Supplementary Figure 2: *Dm*CFL knockdown in the muscle results in progressive loss of both muscle structure and function

**A**

| Genotypes | Class 1 | Class 2 | Class 3 |
|---|---|---|---|
| *control RNAi* vs. *DmCFL RNAi (1)* | **** | ns | **** |
| *control RNAi* vs. *DmCFL RNAi (1)* WT movers | ** | ns | * |
| *control RNAi* vs. *DmCFL RNAi (1)* slow movers | **** | * | **** |
| *DmCFL RNAi (1)* vs. *DmCFL RNAi (1)* WT movers | **** | ns | *** |
| *DmCFL RNAi (1)* vs. *DmCFL RNAi (1)* slow movers | *** | ns | ** |
| *DmCFL RNAi (1)* WT movers vs. *DmCFL RNAi (1)* slow movers | **** | ns | **** |
| | ****p<0.0001 | ***p<0.001 | **p<0.01 *p<0.05 |

**B**

| Larval stages | Class 1 | Class 2 | Class 3 |
|---|---|---|---|
| 1st instar vs. 2nd instar | *** | *** | ns |
| 1st instar vs. early 3rd instar | **** | ** | ** |
| 1st instar vs. late 3rd instar | **** | * | **** |
| 2nd instar vs. early 3rd instar | ns | ns | ** |
| 2nd instar vs. late 3rd instar | *** | * | **** |
| early 3rd instar vs. late 3rd instar | ** | ns | ** |
| | ****p<0.0001 | ***p<0.001 | **p<0.01 *p<0.05 |

**C**

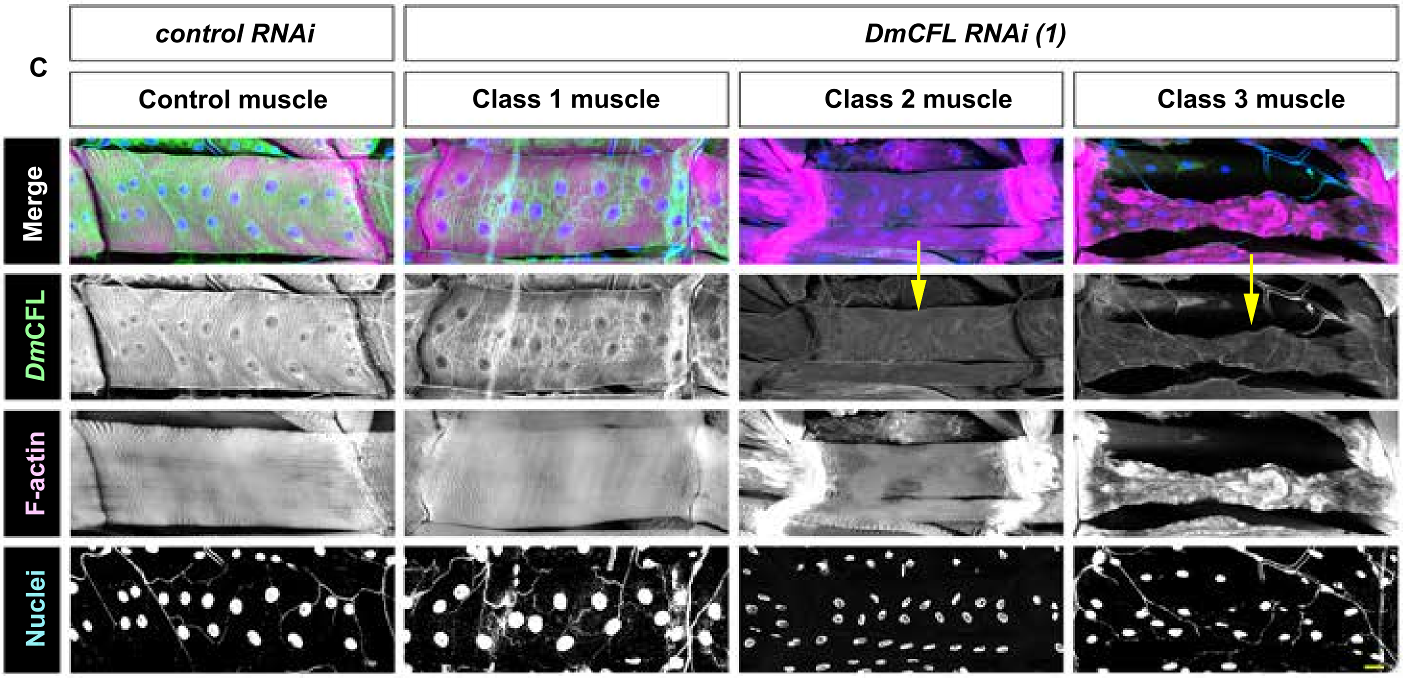

*Balakrishnan et al., Supplementary Figure 2 Legend*

**Supplementary Figure 2: *Dm*CFL knockdown in the muscle results in progressive loss of both muscle structure and function -- Supplement to Figure 3**

A. Statistical analysis of the three muscle classes for the following genotypes: *control RNAi, DmCFL RNAi (1), DmCFL RNAi (1)* WT movers, and *DmCFL RNAi (1)* slow movers. Data shown in Figure 3C. P-values calculated by 2-way ANOVA test.

B. Statistical analysis of the three muscle classes for the $1^{st}$, $2^{nd}$, early $3^{rd}$, and late $3^{rd}$ instars stages of *DmCFL RNAi (1)* larvae. Data shown in Figure 3E. P-values calculated by 2-way ANOVA test.

C. Representative examples of *Dm*CFL expression in $3^{rd}$ instar *DmCFL RNAi (1)* knockdown VL3 larval muscles of each class. *Dm*CFL (green, top row; greyscale row 2), Phalloidin (F-actin, magenta, top row; greyscale row 3), and Hoechst (Nuclei, blue, top row, greyscale row 4). In both *control RNAi* and Class 1 muscles of *DmCFL RNAi (1)* (first and second columns, respectively), *Dm*CFL is localized in identical patterns. In the Class 2 and Class 3 muscles in *DmCFL RNAi* larvae (third and fourth columns respectively), *Dm*CFL is not as highly expressed as in the control cell (yellow arrows). See also S1G. N=200 muscles from 8 larvae of each genotype, repeated in triplicate. Bar, 25µm.

*Balakrishnan et al., Supplementary Figure 3 - Supplement to Figure 4*

### Supplementary Figure 3: *Dm*CFL knockdown results in recruitment of sarcomeric Tropomodulin and Troponin to actin aggregates

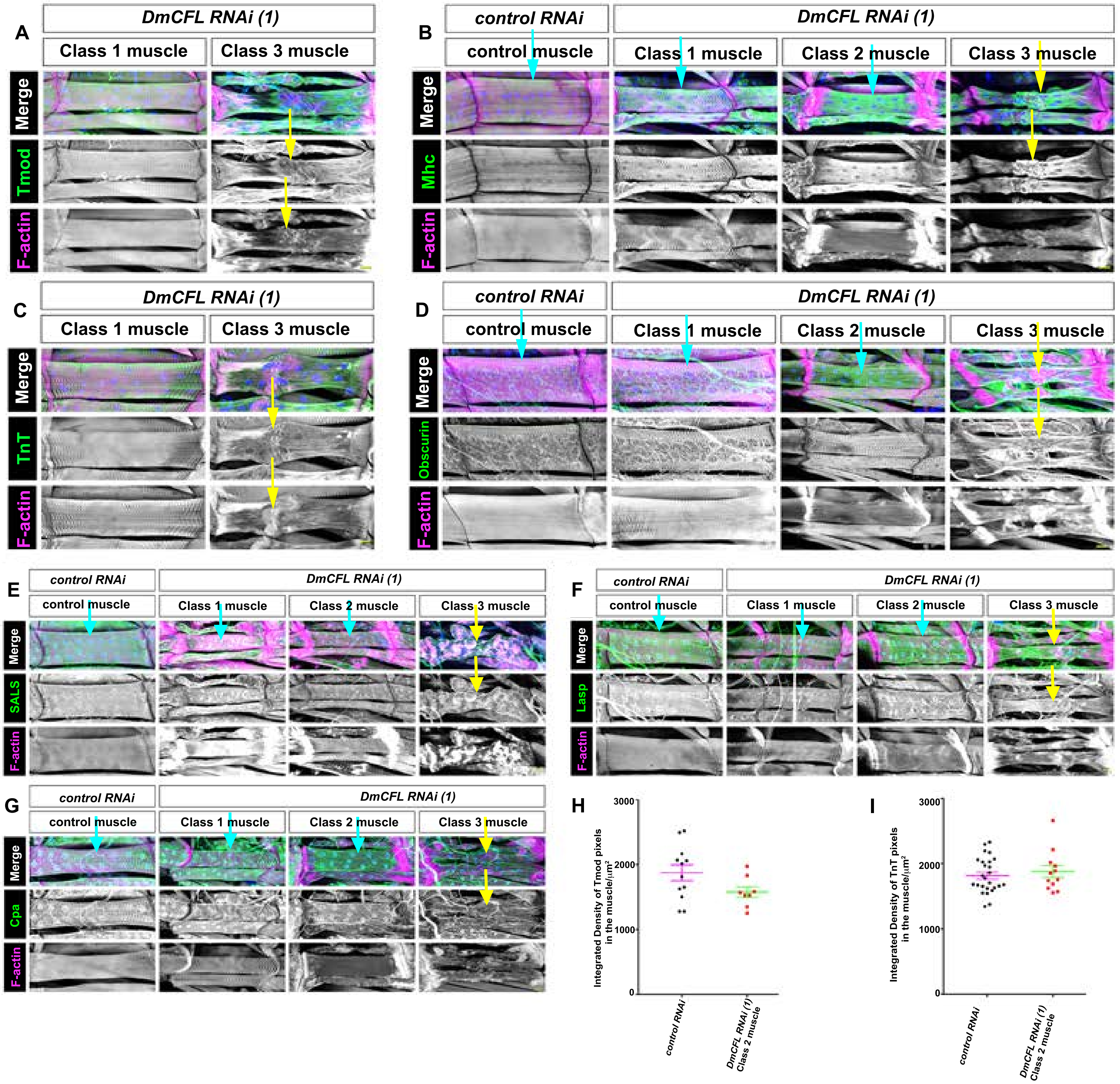

*Balakrishnan et al., Supplementary Figure 3 Legend*

**Supplementary Figure 3: *Dm*CFL knockdown results in recruitment of sarcomeric Tropomodulin and Troponin to actin aggregates – Supplement to Figure 4**

VL3 muscles from *DmCFL RNAi* (1) expressing dissected 3rd instar larvae. N=8 larvae of each genotype, repeated in triplicate for each condition listed below unless stated.

A. Tropomodulin (Tmod, green, top row; greyscale, row 2), Phalloidin (F-actin, magenta, top row; greyscale, row 3), and Hoechst (Nuclei, blue, top row). In Class 1 *DmCFL RNAi (1)* muscles (first column), Tmod localizes in an identical manner to the *control RNAi* muscles (Fig. 4A). In the Class 3 muscles (second column), both actin and Tmod are disorganized.
B. Myosin heavy chain (Mhc) (green, top row; greyscale, row 2), Phalloidin (F-actin, magenta, top row; greyscale, row 3), and Hoechst (Nuclei, blue, top row). In *DmCFL RNAi (1)* Class 1 and 2 muscles (second and third column respectively), Mhc is expressed in an identical pattern as seen in the *control RNAi* muscles (blue arrows). In the Class 3 muscles (fourth column), both actin and Mhc are disorganized (yellow arrows).
C. Troponin T (TnT, green, top row; greyscale, row 2), Phalloidin (F-actin, magenta top row; greyscale, row 3), and Hoechst (Nuclei, blue, top row). In Class 1 *DmCFL RNAi (1)* muscles (first column), TnT localizes in an identical manner to the *control RNAi* muscles (Fig. 4B). In the Class 3 *DmCFL RNAi (1)* muscles (second column), both actin and Tmod are disorganized (yellow arrows).
D. Obscurin (green, top row; greyscale, row 2), Phalloidin (F-actin, magenta, top row; greyscale, row 3), and Hoechst (Nuclei, blue, top row). In the Class 1 and 2 *DmCFL RNAi (1)* muscles (second and third column respectively), Obscurin is expressed in an identical pattern to the *control RNAi* muscle (blue arrows). In the Class 3 *DmCFL RNAi (1)* muscles (fourth column), both actin and Obscurin are disorganized (yellow arrows).
E. SALS (green, top row; greyscale, row 2), Phalloidin (F-actin, magenta, top row; greyscale, row 3), and Hoechst (Nuclei, blue; top row). In the Class 1 and 2 *DmCFL RNAi* (1) muscles

(second and third column respectively), SALS is expressed in an identical pattern to *control RNAi* muscles (blue arrows). In the Class 3 muscle cells (fourth column), both actin and SALS are disorganized (yellow arrows).

F. Lasp (green, top row; greyscale row 2), Phalloidin (F-actin, magenta, top row; greyscale row 3), and Hoechst (Nuclei, blue, top row). In the Class 1 and 2 *DmCFL RNAi (1)* muscles (second and third column respectively), Lasp is expressed in an identical pattern to *control RNAi* muscles (blue arrows). In the Class 3 *DmCFL RNAi (1)* muscles (fourth column), both actin and Lasp are disorganized (yellow arrows).

G. Capping protein a (Cpa) (green, top row; greyscale, row 2), Phalloidin (F-actin, magenta, top row; greyscale, row 3), and Hoechst (Nuclei, blue, top row). In the Class 1 and 2 *DmCFL RNAi (1)* muscles of (second and third column respectively), Cpa is expressed in an identical pattern to *control RNAi* muscles (blue arrows). In the Class 3 *DmCFL RNAi (1)* muscles (fourth column), both actin and Cpa are disorganized (yellow arrows).

H. Integrated density of Tropomodulin (Tmod) pixels per $\mu m^2$ of muscle cell area. Pixel intensity was calculated from *control RNAi* and Class 2 *DmCFL RNAi (1)* muscles. Example images are shown in Figure 4A. There is no significant difference in Tmod pixel intensity between Class 2 *DmCFL RNAi (1)* muscles and control muscles (ns, p=0.087). Error bars, mean±SEM. N ≥10 muscles from at least 5 different larvae, repeated in duplicate.

I. Integrated density of Troponin T (TnT) pixels per $\mu m^2$ of muscle cell area. Pixel intensity was calculated from *control RNAi* and Class 2 *DmCFL RNAi (1)* muscles. Example images are shown in Fig. 4B. There is no significant difference in TnT pixel intensity between Class 2 *DmCFL RNAi (1)* muscles and control muscles (ns, p=0.2417). Error bars, mean±SEM. N ≥10 muscles from at least 5 different larvae, repeated in duplicate.

Scale Bar, 25μm.

### Supplementary Figure 4: *Dm*CFL knockdown leads to fewer sarcomeres being added in growing muscle cells

**A**

| Genotypes | p-value |
|---|---|
| *control RNAi vs. Class 1 muscle DmCFL RNAi (1)* | * |
| *control RNAi vs. Class 2 muscle DmCFL RNAi (1)* | **** |
| *Class 1 muscle DmCFL RNAi (1) vs. Class 2 muscle DmCFL RNAi (1)* | * |
| * p<0.05 | ****p<0.0001 |

*Balakrishnan et al., Supplementary Figure 4 Legend*

**Supplementary Figure 4: *Dm*CFL knockdown leads to fewer sarcomeres being added in growing muscle cells. – Supplement to Figure 5**

A. Statistical analysis performed on sarcomere number in VL muscles of the same length between the listed genotypes. There was significant difference in the number of sarcomeres in Class 2 *DmCFL RNAi (1)* muscles in comparison to both Class 1 *DmCFL RNAi (1)* muscles and muscles from *control RNAi*-expressing larvae. P-values calculated by 1-way ANOVA test.

*Balakrishnan et al., Supplementary Figure 5 - Supplement to Figure 6*

### Supplementary Figure 5: Modulating proteasome activity affects the progression of structural and functional changes in *Dm*CFL-knockdown larval muscle cells

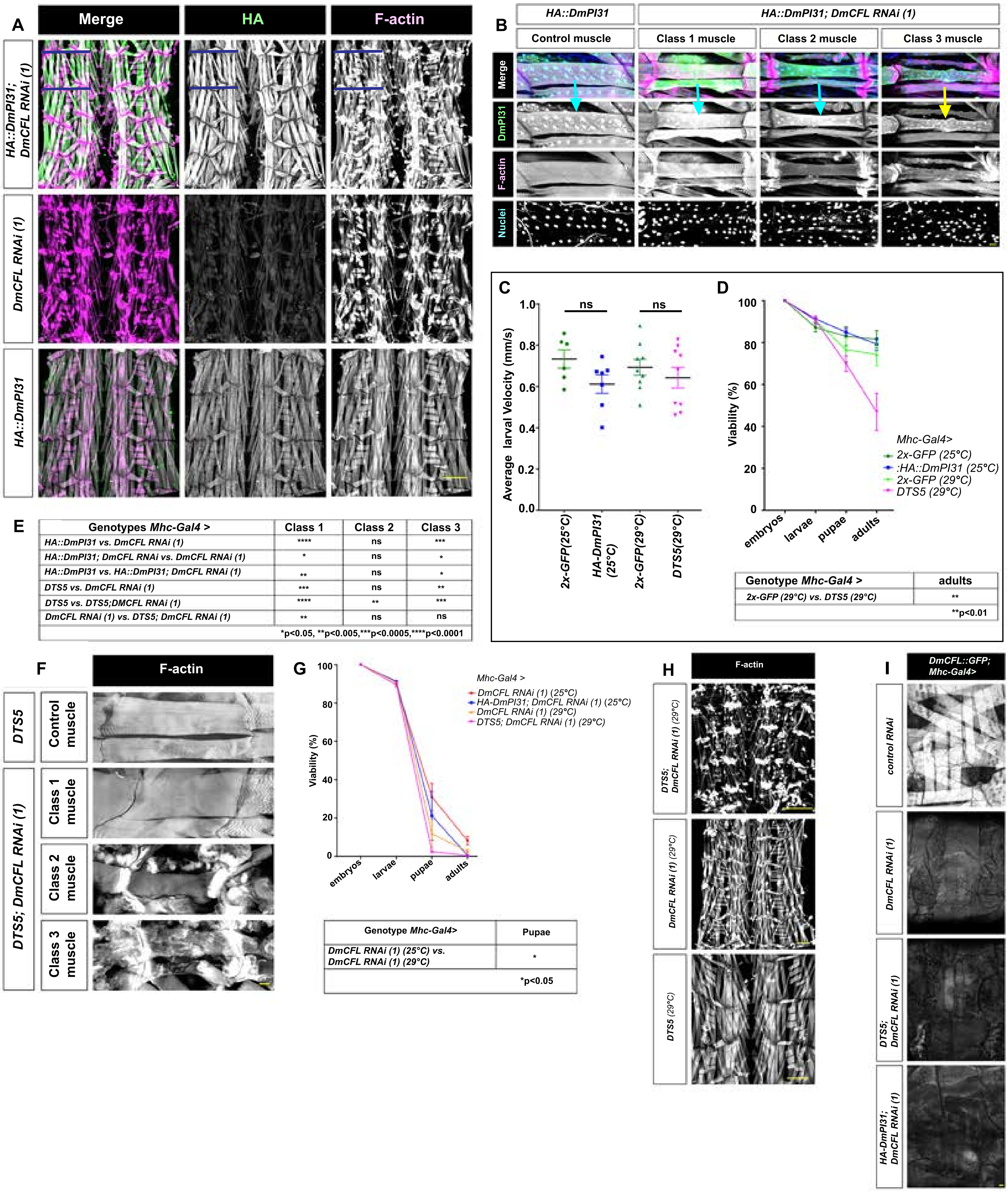


E

| Genotypes *Mhc-Gal4 >* | Class 1 | Class 2 | Class 3 |
|---|---|---|---|
| HA::DmPI31 vs. DmCFL RNAi (1) | **** | ns | *** |
| HA::DmPI31; DmCFL RNAi vs. DmCFL RNAi (1) | * | ns | * |
| HA::DmPI31 vs. HA::DmPI31; DmCFL RNAi (1) | ** | ns | * |
| DTS5 vs. DmCFL RNAi (1) | *** | ns | ** |
| DTS5 vs. DTS5;DMCFL RNAi (1) | **** | ** | *** |
| DmCFL RNAi (1) vs. DTS5; DmCFL RNAi (1) | ** | ns | ns |
| | *p<0.05, **p<0.005,***p<0.0005,****p<0.0001 | | |

D

| Genotype *Mhc-Gal4 >* | adults |
|---|---|
| 2x-GFP (29°C) vs. DTS5 (29°C) | ** |
| | **p<0.01 |

G

| Genotype *Mhc-Gal4>* | Pupae |
|---|---|
| DmCFL RNAi (1) (25°C) vs. DmCFL RNAi (1) (29°C) | * |
| | *p<0.05 |

*Balakrishnan et al., Supplementary Figure 5 Legend*

**Supplementary Figure 5: Modulating proteasome activity affects the progression of structural and functional changes in *Dm*CFL-knockdown larval muscle cells. – Supplement to Figure 6**

A. Comparison of the musculature of four abdominal segments of filleted 3rd instar larvae of the listed genotypes. HA (green, first column; greyscale, column 2) and Phalloidin (F-actin, magenta, first column; greyscale, column 3). Blue lines indicate one larval hemi-segment. N=8 larvae of each genotype, repeated in triplicate.

B. VL3 muscle cells from dissected 3rd instar larvae expressing *HA::DmPI31* or *HA::DmPI31; DmCFL RNAi (1)* . HA (green, top row; greyscale, row 2), Phalloidin (F-actin, magenta, top row; greyscale, row 3), and Hoechst (Nuclei, blue, top row; greyscale, row 4). In the Class 1 and 2 *HA::DmPI31; DmCFL RNAi (1)* muscles (second and third columns respectively), HA::DmPI31 localizes in same pattern seen in the *control RNAi* muscles (blue arrows). As the muscle architecture is disorganized in Class 3 muscles (yellow arrow, fourth column), DmPI31's localization is also disorganized. N=8 larvae of each genotype, repeated in triplicate.

C. Crawling velocities of 3rd instar larvae with *Mhc-Gal4* driving expression of the listed UAS-constructs. There is no significant difference in the average velocity between the control and *HA::DmPI31* or *DTS5* expressed in control larvae. Error bars mean ±SEM. N=8 larvae per genotype, repeated twice.

D. Viability of *HA::DmPI31* and *DTS5* expressing flies compared to controls raised at identical temperatures. Muscle-specific expression of DTS5 decreases the percentage of pupae and adults, indicating that inhibiting the proteasome in the larval musculature can affect the overall viability of the organism. $^{**}p<0.01$. P values calculated by 2-way ANOVA.

N=100 embryos of each genotype per experiment tracked over the different developmental stages. Experiments were repeated 2 times. Error bars, mean ±SEM.

E. Statistical analysis of the three classes of muscles between the different genotypes (Figure 6G and 6I). P-values calculated by 2-way ANOVA test.

F. VL3 muscle cells from dissected 3rd instar larvae of the listed genotypes. F-actin (greyscale). N=8 larvae of each genotype, repeated in triplicate.

G. Viability of flies expressing *HA::DmPI31* or *DTS5* with *DmCFL RNAi (1)*. No significant decrease in the viability of the organism between the different genotypes was detected. While there is a small significant difference in the percentage of pupae formed in *DmCFL RNAi (1)* 25°C vs *DmCFL RNAi (1)* 29°C, it did not affect the number that adult flies that eclose. $^{*}p<0.05$. P values calculated by 2-way ANOVA. N=100 embryos of each genotype per experiment tracked over the different developmental stages. Experiments were repeated twice. Error bars, mean±SEM.

H. Comparison of the musculature of four abdominal segments of filleted 3rd instar larvae of the listed genotypes. Phallodin (F-actin, greyscale). Reduction of proteasome activity enhances the sarcomere phenotypes. N=8 larvae of each genotype, repeated in triplicate.

I. *Dm*CFL::GFP (greyscale) expressing muscles at 3rd instar with the different proteasome constructs. *Dm*CFL levels are not affected by either activation or inhibition of the proteasome. N= 4 larvae of each genotype, repeated twice.

Scale Bar: 250μm (A,H), 25μm (B, F, I).

*Balakrishnan et al., Supplemental Figure 6 - Supplement to Figure 7*

### Supplementary Figure 6: *In vivo* and *in vitro* analyses identify phenotypic and biochemical consequences of a specific patient *CFL2* mutation

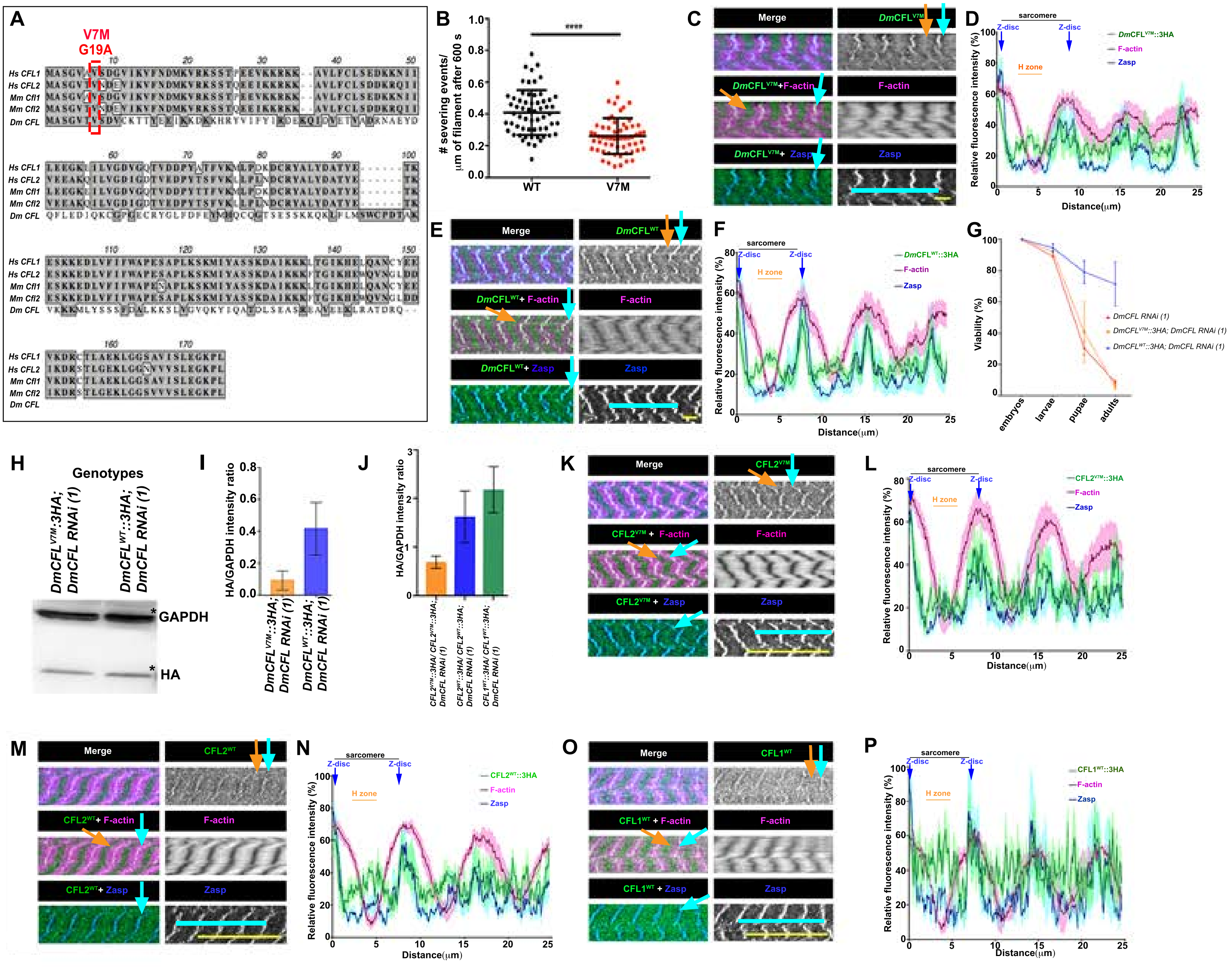


Q

| Genotypes *Dmef2-Gal4>* | Muscle Classes | | Velocities |
|---|---|---|---|
| | Class 1 | Class 3 | |
| *DmCFL RNAi (1) & control RNAi* | **** | **** | **** |
| *DmCFL RNAi (1) & CFL2$^{V7M}$::3HA; DmCFL RNAi (1)* | ns | ns | * |
| *DmCFL RNAi (1) & CFL2$^{WT}$::3HA; DmCFL RNAi (1)* | *** | *** | **** |
| *DmCFL RNAi (1) & CFL1$^{WT}$::3HA; DmCFL RNAi (1)* | *** | *** | **** |
| *DmCFL RNAi (1) & CFL2$^{V7M}$::3HA/ CFL2$^{V7M}$::3HA; DmCFL RNAi (1)* | *** | *** | *** |
| *DmCFL RNAi (1) & CFL2$^{WT}$::3HA/CFL2$^{WT}$::3HA; DmCFL RNAi (1)* | **** | **** | **** |
| *DmCFL RNAi (1) & CFL1$^{WT}$::3HA/CFL1$^{WT}$::3HA; DmCFL RNAi (1)* | **** | **** | **** |
| | *p<0.05 | ***p<0.001 | ****p<0.0001 |

**Supplementary Figure 6: *In vivo* and *in vitro* analyses identify phenotypic and biochemical consequences of a specific patient *CFL2* mutation. –Supplement to Figure 7**

A. Sequence alignment of entire *Homo sapiens* Cofilin-1 (CFL1) and Cofilin-2 (CFL2), *Mus musculus* CFL1 and CFL2, and *Drosophila melanogaster* 's *Dm*CFL. Grey regions denote regions sharing homology.

B. Analysis of actin filament severing activity after 600 seconds. Each data point is the total number of severing events that occurred during reaction (600 seconds) per micron of F-actin, compiled from three independent experiments (20 actin filaments each). ****p<0.0001. P values calculated by Unpaired t-test with Welch's Correction factor. Error bars, mean ±SD.

C. Single Z slice of a portion of a VL3 muscle expressing *Dm*CFL$^{V7M}$::3HA. HA (*Dm*CFL $^{V7M}$::3HA, green), Phalloidin (F-actin, magenta), and Zasp (Z-disc, blue). *Dm*CFL$^{V7M}$ co-localizes with F-actin and Zasp (blue arrow). *Dm*CFL$^{V7M}$ also localizes to the F-actin free region of the sarcomere i.e. H-zone (orange arrow). *Dm*CFL $^{V7M}$ localizes to the same regions of the muscle as endogenous *Dm*CFL (Figure 1G). Blue line is example of 4 sarcomeres quantified in (D). N=5 larvae, repeated twice.

D. Relative average fluorescence intensities of *Dm*CFL$^{V7M}$ (green), F-actin (magenta), and Zasp (blue) across the length of 4 sarcomeres (average ± SEM). *Dm*CFL$^{V7M}$ localizes to the Z disc of the sarcomere (blue arrow) as well as the F-actin free region of the sarcomere, i.e. the H-zone (orange line). N = 9 measurements of 4 sarcomeres [blue line in (C)] from 8 different muscles of 5 larvae.

E. Single Z slice of a portion of a VL3 muscle expressing *Dm*CFL$^{WT}$::3HA. HA (*Dm*CFL $^{WT}$::3HA, green), Phalloidin (F-actin, magenta), and Zasp (Z-disc, blue). *Dm*CFL$^{WT}$ co-localizes with F-actin and Zasp (blue arrow). *Dm*CFL$^{WT}$ also localizes to the F-actin free region of the sarcomere i.e. H-zone (orange arrow). *Dm*CFL$^{WT}$ localizes to the same regions of the muscle as endogenous *Dm*CFL (Figure 1G). Blue line is example of 4 sarcomeres quantified in (F). N=5 larvae, repeated twice.

F. Relative average fluorescence intensities of *Dm*CFL$^{WT}$ (green), F-actin (magenta), and Zasp (blue) across the length of 4 sarcomeres (average $\pm$ SEM). *Dm*CFL$^{WT}$ localizes to the Z disc of the sarcomere (blue arrow) as well as the F-actin free region of the sarcomere, i.e. the H-zone (orange line). N = 9 measurements of 4 sarcomeres [blue line in (E)] from 8 different muscles of 5 larvae.

G. Viability of *DmCFL RNAi (1)* flies co-expressing *DmCFL*$^{V7M}$ or Dm*CFL*$^{WT}$ using *Mhc-Gal4*. Expression of *DmCFL*$^{WT}$ with *DmCFL RNAi (1)* rescues viability of the organism by increasing the percentage of pupae and adults. Expression of *DmCFL*$^{V7M}$ with *DmCFL RNAi (1)* did not significantly rescue viability of the organism. N=100 embryos of each genotype per experiment tracked over the different developmental stages. Experiments were repeated twice. P-values calculated by 2-way ANOVA test. Statistical analysis is shown in Supplementary Figure 1F. Error bars, mean±SEM.

H. Example immunoblot depicting HA levels of the listed proteins in 3$^{rd}$ instar larval carcasses (consisting primarily of muscle). GAPDH is used as an internal loading control for all the samples. Quantification of multiple repeats shown in (I).

I. Quantification of HA protein expression of the listed constructs. HA levels were normalized to each sample's GAPDH levels. 20 carcasses were analyzed in triplicate for each genotype. Error bars, mean ±SEM.

J. Quantification of HA protein expression levels of listed constructs. HA levels were normalized to each sample's GAPDH levels. 20 carcasses were analyzed in triplicate for each genotype. Expressing two copies of human *CFL2*$^{WT}$ and *CFL1*$^{WT}$ in a *Dm*CFL depleted muscle increase the HA levels. This contributed to the better levels of rescue seen in these larvae. Error bars, mean ±SEM.

K. Single Z slice of a portion of a VL3 muscle expressing *Hs*Cofilin-2$^{V7M}$ (CFL2$^{V7M}$)::3HA. \ HA (CFL2$^{V7M}$::3HA, green), Phalloidin (F-actin, magenta), and Zasp (Z-disc, blue). CFL2$^{V7M}$ co-localizes with F-actin and Zasp at the Z disc (blue arrow). CFL2$^{V7M}$ also localizes to the F-actin free region of the sarcomere i.e. H-zone (orange arrow). CFL2$^{V7M}$ localizes to the same regions of the muscle as endogenous *Dm*CFL (Figure 1G). Blue line is example of 4 sarcomeres quantified in (L). N=5 larvae, repeated twice.

L. Relative average fluorescence intensities of *Hs*Cofilin-2$^{V7M}$ (CFL2$^{V7M}$,green), F-actin (magenta), and Zasp (blue) across the length of 4 sarcomeres (average ± SEM). CFL2$^{V7M}$ localizes to the Z disc of the sarcomere (blue arrow) as well as the F-actin free region of the sarcomere, i.e. the H-zone (orange line). N = 9 measurements of 4 sarcomeres [blue line in (K)] from 8 different muscles of 5 larvae.

M. Single Z slice of a portion of a VL3 muscle expressing *Hs*Cofilin-2$^{WT}$::3HA (CFL2$^{WT}$). HA (CFL2$^{WT}$::3HA, green), Phalloidin (F-actin, magenta), and Zasp (Z-disc, blue). CFL2$^{WT}$ co-localizes with F-actin and Zasp at the Z disc (blue arrow). CFL2$^{WT}$ also localizes to the F-actin free region of the sarcomere i.e. H-zone (orange arrow). CFL2$^{WT}$ localizes to the same regions of the muscle as endogenous *Dm*CFL (Figure 1G). Blue line is example of 4 sarcomeres quantified in (N). N=5 larvae, repeated twice.

N. Relative average fluorescence intensities of *Hs*Cofilin-2$^{WT}$(CFL2$^{WT}$,green), F-actin (magenta), and Zasp (blue) across the length of 4 sarcomeres (average ± SEM). CFL2$^{WT}$ localizes to the Z disc of the sarcomere (blue arrow) as well as the F-actin free region of the sarcomere, i.e. the H-zone (orange line). N = 9 measurements of 4 sarcomeres [blue line in (M)] from 8 different muscles of 5 larvae.

*Balakrishnan et al., Supplementary Figure 6 Legend*

O. Single Z slice of a portion of a VL3 muscle expressing *Hs*Cofilin-1$^{WT}$(CFL1$^{WT}$) ::3HA. HA (CFL1$^{WT}$, green), Phalloidin (F-actin, magenta), and Zasp (Z-disc, blue). CFL1$^{WT}$ co-localizes with F-actin and Zasp at the Z disc (blue arrow). CFL1$^{WT}$ also localizes to the F-actin free region of the sarcomere i.e. H-zone (orange arrow). CFL1$^{WT}$ localizes to the same regions of the muscle as endogenous *Dm*CFL (Figure 1G). Blue line is example of 4 sarcomeres quantified in (P). N=5 larvae, repeated twice.

P. Relative average fluorescence intensities of *Hs*Cofilin-1$^{WT}$(CFL1$^{WT}$, green), F-actin (magenta), and Zasp (blue) across the length of 4 sarcomeres (average ± SEM). CFL1$^{WT}$ localizes to the Z disc of the sarcomere (blue arrow) as well as the F-actin free region of the sarcomere the H-zone (orange arrow). N = 9 measurements of 4 sarcomeres [blue line (O)] from 8 different muscles of 5 larvae.

Q. Statistical analysis on the velocities and classes of muscles between the listed genotypes. There was no significant difference in the viability of the genotypes at the embryo and larval stages, yet significant differences at subsequent stages. P-values calculated by 2-way ANOVA test for muscle classes and by One way-ANOVA for the larval velocities.

Scale Bar: 25µm (C, E, K, M,O)
